# Decomposing the modulation of interactions between neuronal populations

**DOI:** 10.64898/2026.05.14.725145

**Authors:** Marco Celotto, J. Samuel Sooter, Sofie Ährlund-Richter, Kyle R. Jenks, Mriganka Sur, Stefano Panzeri

**Affiliations:** Picower Institute for Learning and Memory, Massachusetts Institute of Technology, USA; Institute of Neural Information Processing, University Medical Center Hamburg-Eppendorf (UKE), Hamburg, Germany; Department of Physics, University of Arkansas, USA

## Abstract

Identifying, through analysis of large-scale neural recordings, subpopulations of neurons interacting across brain areas is key for understanding across-brain communication with cellular resolution. The communication subspace (CS) framework addresses this need by capturing additive interactions between subspaces of source and target neural populations. However, no current method identifies interpretable components of how a measured, multivariate modulator - such as behavioral state or the activity of another population - reshapes these interactions. Here we extend the CS framework by using a bilinear perturbation of reduced-rank regression to parameterize modulation as a low-rank tensor. We derive MICs - multiplicative interaction channels - a method to identify triplets made of a source and a target activity pattern and a modulator of their interaction. We develop a hierarchical fitting pipeline and provide a closed-form decomposition that quantifies whether modulation reshapes the modulator-averaged baseline interaction, recruits private dimensions of one population, or opens new interactions. In simulations, MICs reliably recovers the presence and geometry of ground-truth modulation even in the high-dimensional, low-sample regime. Applied to a benchmark macaque dataset, MICs recovers the geometry and functional form of the stimulus modulation of interactions between primary and secondary visual cortices. Applied to simultaneous calcium imaging of prefrontal axons and interneurons in mouse visual cortex, MICs reveals that behavioral state asymmetrically modulates top-down interactions, reconfiguring the prefrontal projection patterns that interact with a stable set of visual interneurons. MICs provides a data-efficient, interpretable account of how measured modulators reshape the interactions between neural populations. Code is available at https://github.com/mcelotto/multiplicative_interaction_channels.

## 1 Introduction

Brain computations supporting cognitive functions arise from interactions between multiple brain regions [1, 2]. Inter-area interactions are implemented in the brain with cellular-level resolution [3–5], as specific subpopulations of neurons dynamically interacting with other subpopulations. As modern recording technologies access hundreds to thousands of neurons across multiple regions [6–8], a fundamental computational challenge is to develop analysis tools that can map these interactions from simultaneous recordings of many neurons and work even in the under-sampled regime typical of brain data [9]. Classic pairwise approaches based on correlation [10] or the Wiener-Granger principle [11–14] were developed for scalar variables or small populations are impractical for high-dimensional recordings [15]. The *Communication Subspace* (CS; see Glossary A) [16] has emerged as a powerful framework to address this challenge. It uses Reduced-rank regression (RRR) to constrain the regression matrix relating two populations to low-rank to identify low-dimensional patterns of activity of one population that co-vary with low-dimensional activity patterns of the other, excluding variability private to each population [17]. The low-rank constraint dramatically reduces the number of free parameters, achieving data-efficiency and scalability. CS has become a popular tool to study inter-areal brain interactions with cellular resolution [18–22].

However, interactions between neural populations can change with cognitive state, movement parameters, task context, and activity of other brain regions. For example, attention *modulates* top-down interactions between higher cortical areas and sensory cortex [23, 24]. Yet the population-level structure of these modulations remains unknown. Modulation effects of a modulator on the relationship between other variables are usually described as multiplicative - rather than additive [25–28]. Moreover, modulatory variables can themselves be high-dimensional, such as multiple movement parameters [29] or the activity of other populations [30]. Despite the importance of modulatory interactions, CS analysis captures additive pairwise interactions between populations, but not the multiplicative modulation of their interaction by a third, potentially high-dimensional variable.

Here, we derive Multiplicative Interactions Channels (*MICs*), a method extending communication subspaces to capture how a high-dimensional modulator changes interactions between a source and a target population (1). We model the modulator-induced perturbation of the source-target mapping as a low-rank three-way tensor and parameterize it via a Canonical Polyadic (CP) decomposition [31–33]. Low-rank tensor decompositions can be robustly applied to neural population data [34–36] and to regression with multivariate covariates [37–39]. We thus parameterize the perturbation as a sum of rank-1 *multiplicative interaction channels*, each pairing one modulator pattern with the source-target interaction pattern it gates. We describe a hierarchical data-fitting procedure with cross-validated rank and regularization selection, and show that the geometry of each MIC relative to the modulator-averaged, baseline communication subspace can be decomposed into four interpretable components. We next validate MIC on simulated data. We then apply it to a public macaque V1-V2 benchmark dataset, recovering the expected stimulus dependence of the communication subspace with a single model and resolving its form and geometry, and to new simultaneous recordings of prefrontal axonal projections and visual cortical interneurons in mice.

## 2 Related work

Several methods model dynamically modulated interactions within or between neuronal populations, but none quantifies how a measured modulator reshapes the mapping between a source and a target population and describes the geometry of this modulation. They differ from MICs along at least one of two axes. First, *how the modulation enters*. In multi-region switching dynamical systems [40, 41], the interaction between regions changes with an inferred discrete state. In sequential autoencoders of inter-regional communication [42], messages between regions are inferred, and a third region can change the interaction only through the network’s nonlinearity. In decomposed dynamical systems [43, 44], the dynamics are a combination of basis matrices with latent coefficients, from which behavior can be predicted [44]. These models can report that an interaction changed, but a measured variable never enters as an explicit modulator that the interaction depends on. Second, *what is modulated*. Low-rank tensor autoregressions [45, 46], behavior-constrained decomposed dynamics [44], and conditionally linear dynamical systems [47] modulate the recurrent dynamics of a single population. Several of these methods [43–47] share with MICs a decomposition into basis matrices. MICs differ in that the coefficients are functions of a measured modulator, the matrices map source to target, and each channel admits a geometric decomposition relative to the baseline interaction.

## 3 Mathematical derivation of multiplicative interaction channels

### 3.1 Background: the Communication Subspace (CS) framework

We consider simultaneous observations of three possibly high-dimensional variables: a *source* 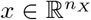, a *target* 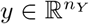, and a *modulator* 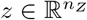 . For example, *x* and *y* could represent the activity of two neural populations, and *z* could be a set of behavioral variables (e.g., running speed, pupil diameter), or the activity of a third modulatory population. Given *N* simultaneous observations, such as time points in a multivariate time series, we build data matrices 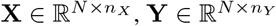, and 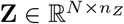. The CS framework [16] identifies components of source activity linearly related to other specific components of target activity. It models the target as a low-rank linear function of the source via RRR:

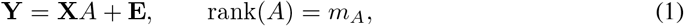

where 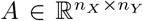 and **E** denotes residual variability. The rank constraint implies that *A* can be factorized as 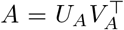, with 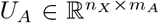 and 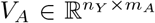. The columns of *U*_*A*_ and *V*_*A*_ define, respectively, the *m*_*A*_ source and target patterns that co-vary with each other; patterns outside col(*U*_*A*_) and col(*V*_*A*_) are private to each population [16]. The optimal *A* is obtained by ridge-RRR [48, 17] (Appendix B.1). We refer to col(*U*_*A*_) and col(*V*_*A*_) as the source and target *additive interaction subspaces* (AIS), rather than “communication subspaces” [16], as RRR at zero lag captures additive co-variation, not directed transmission [11, 12, 17], and source/target swapping does not, on its own, establish directionality [18]. Next, we will consider how to extend the CS framework beyond additive interactions.

### 3.2 Extending CS to multiplicative interactions beyond Additive Interaction Subspaces (AIS)

AIS capture *additive*, linear relationships between two variables. However, many interactions in neuroscience are hypothesized to be *modulated* by additional variables. We thus ask how the modulator *Z* changes the linear mapping from source to target (Fig. 1a). We model the instantaneous effective interaction operator as a *z*-dependent perturbation of the baseline mapping:

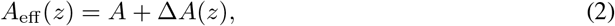

**Figure 1:**
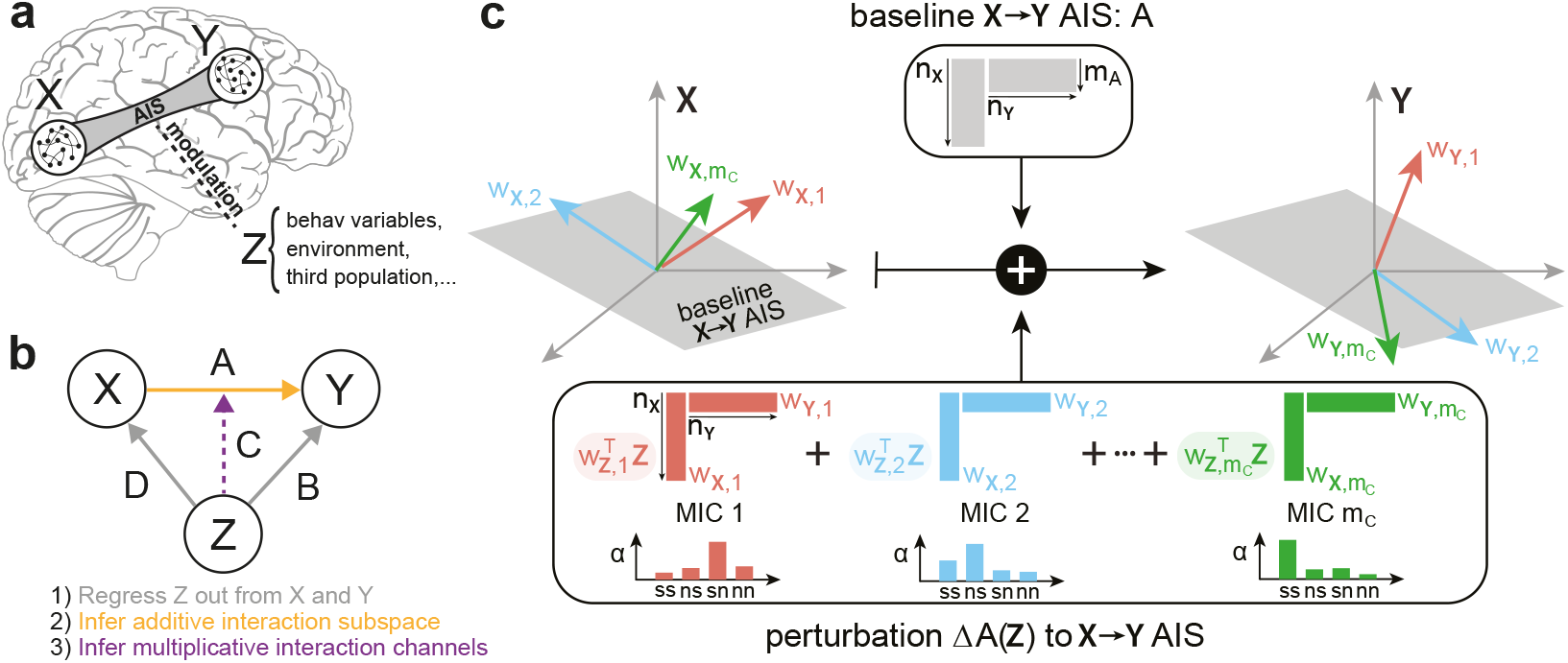
Sketch of MICs. (a) MICs quantify how the AIS between a multivariate source *X* and target *Y* changes as a function of a modulator *Z*. (b) Schematic of the modulation-channel fitting pipeline. (c) The perturbation Δ*A*(*z*) to the baseline *X* → *Y* mapping *A* is parameterized as a low-rank tensor: low-dimensional components of *X* and *Z* interact bilinearly to drive specific activity patterns in *Y* . Each channel’s source and target directions need not lie within the baseline AIS; their geometry relative to it is summarized by four indices (*α*_*ss*_, *α*_*ns*_, *α*_*sn*_, *α*_*nn*_; see Sec. 3.4).

where *A* is the baseline AIS and Δ*A*(*z*) captures the *z*-dependent modulation. We assume Δ*A*(*z*) to be linear in *z*, so that its action on the source is bilinear in *x* and *z*, with linearity in *x* at fixed *z* inherited from RRR. Bilinearity is deliberate: together with the low-rank parameterization below, it is what makes MICs sample-efficient (Section 4.1). We thus parameterize Δ*A*(*z*) as a sum of *m*_*C*_ rank-1 *multiplicative interaction channels*^1^:

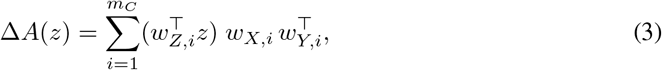

where for each channel *i*, the weight vectors 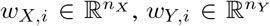, and 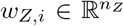 define source, target, and modulator axes, respectively. Intuitively, MIC *i* captures the fact that when the modulator aligns with *w*_*Z,i*_ (i.e., the 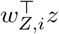 *score* is large), the effective mapping from source to target is perturbed by the rank-1 component 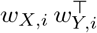, scaled proportionally to 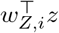. This is a bilinear (multiplicative) interaction between source and modulator, analogous to an interaction term in a regression model, but operating in the high-dimensional setting where source, target, and modulator may each be high-dimensional (see Appendix B.2).

In matrix form, defining *W*_*X*_, *W*_*Z*_, *W*_*Y*_ as the factor matrices whose columns are {*w*_*X,i*_ }, {*w*_*Z,i*_ }, {*w*_*Y,i*_} respectively, and letting ⊙ denote the element-wise (Hadamard) product, the modulation term across all *N* observations can be written compactly as:

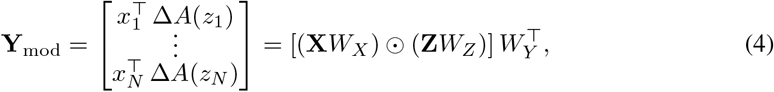

where 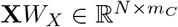 and 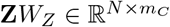are the source and modulator scores (projections onto the respective axes), and their element-wise product captures the bilinear interaction. Mathematically, the collection 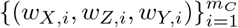 defines a rank-*m*_*C*_ CP decomposition [31–33] of a three-way tensor 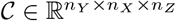.

Beyond the baseline interaction *A* and the modulation tensor *C*, the modulator may also exert direct additive effects on the target and on the source. We capture these with two additional low-rank linear terms: 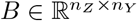(rank *m*_*B*_) for the direct effect *Z* → *Y*, and 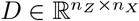 (rank *m*_*D*_) for the effect *Z* → *X*. Letting **X**^*′*^ = **X** − **Z***D* denote the *residualized source* (the component of source variability that is not linearly predictable from the modulator), the complete model is:

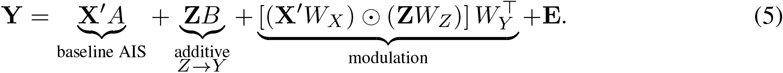

The residualized source **X**^*′*^ enters both the baseline interaction and the modulation terms. In Eq. 5, *A* and *B* capture the first-order effects of *X* and *Z* on *Y*, the modulation term captures their second-order *X* × *Z* interaction. The modulation term is a specific instance of CP-rank-constrained tensor-on-tensor regression [38], with the predictor restricted to the rank-1 outer product *x*_*n*_ ⊗ *z*_*n*_ of two distinct observed variables. It is this restriction, rather than the CP structure of *C* itself, that carries the bilinear-interaction form. The full MICs model (Eq. 5) embeds this term within an additive RRR framework with residualized source, a configuration not addressed in [38] (see Appendix B.2).

#### The baseline AIS operator A

We assume all variables *X, Y*, and *Z* are zero-centered at the stage of model fitting (see Section 3.3). The expected value of the modulatory perturbation is thus 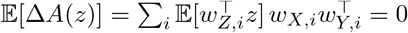. Thus, *A* represents the average effective AIS E[*A*_eff_ (*z*)] = *A*. MICs capture how each observation deviates from this average, depending on *z*.

### 3.3 Hierarchical model fitting and modulatory explained variance

Fitting all parameters of Eq. 5 jointly is computationally challenging due to the interaction between the additive and multiplicative terms. Instead, we adopt a hierarchical procedure that sequentially isolates each component (Algorithm 1; Appendix B.6; Fig. 1b).

#### Step 1: Regressing out additive effects of *Z*

We first fit the additive effects of the modulator on both the target and the source via ridge-RRR:

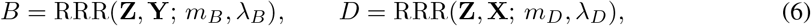

where RRR(·,· ; *m, λ*) denotes ridge-RRR with rank *m* and penalty *λ* (see Appendix B.1), and compute residuals **X**^*′*^ = **X** − **Z***D*, **Y**^*′*^ = **Y** − **Z***B*. This removes variance in both *X* and *Y* that is linearly attributable to **Z** alone. Regressing *Z* out of *X* (via *D*) ensures that downstream terms capture *X*-*Y* relationship not confounded by *Z* independently driving both *X* and *Y* (see Sec. 4.1).

#### Step 2: Fitting baseline interaction

Given the residuals, we fit the baseline interaction matrix as:

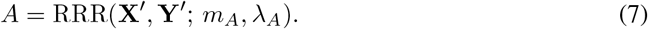

#### Step 3: Fitting MICs

We fit the modulation tensor to the residuals of the baseline interaction model, **Y**^*′′*^ = **Y**^*′*^ − **X**^*′*^*A*, by minimizing the regularized loss:

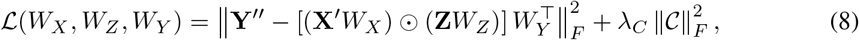

where 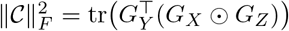 is the squared Frobenius norm of the tensor expressed in terms of the Gram matrices 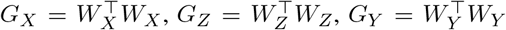 (see Appendix B.4). We minimize Eq. 8 via alternating least squares (ALS), cyclically updating each factor matrix while holding the other two fixed (Algorithm 2, [33, 38]). Each sub-problem reduces to a regularized linear system; closed-form update equations and computational details are provided in Appendix B.5. Eq. 8 is non-convex, so ALS is not guaranteed to reach the global optimum.

#### Rank and regularization selection

Each rank (*m*_*A*_, *m*_*B*_, *m*_*C*_, *m*_*D*_) and ridge penalty (*λ*_*A*_, *λ*_*B*_, *λ*_*C*_, *λ*_*D*_) is selected via *K*-fold cross-validation. For a given rank, the ridge penalty is optimized using Bayesian optimization [50]. The rank is selected as the smallest value whose cross-validated performance is within one SE of the maximum [16, 51]. The same cross-validation folds are used across all steps preventing information leakage between hierarchical stages (Algorithm 1).

#### Modulatory explained variance

Our primary measure of modulatory interaction strength is the *modulatory explained variance* (EV_mod_): the cross-validated *R*^2^ increment of the full model (Eq. 5) over the joint additive model **X**^*′*^*A* + **Z***B*, averaged over folds (Appendix B.6). A positive EV_mod_ indicates that the bilinear *X*-*Z* interaction explains target variance beyond the combination of additive terms. Analogously, EV_AIS_ is the increment from adding **X**^*′*^*A* over the modulator-only prediction **Z***B*, yielding a variance partitioning of **Y** along the hierarchical fit.

### 3.4 Geometric decomposition of MICs

A MIC could modulate interaction geometry by rehaping baseline source-target interactions, engageing private source or target dimensions, or opening up new interaction pathways. We formalize these possibilities by decomposing each channel relative to the baseline AIS.

Let 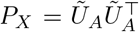 and 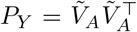 be orthogonal projectors onto the baseline source and target AIS, respectively, where *Ũ*_*A*_ and 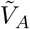 are orthonormal bases for col(*U*_*A*_) and col(*V*_*A*_). Using *I* = *P* + (*I* − *P*) on both the source and target sides, we decompose each channel’s rank-1 perturbation 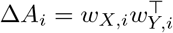 into four orthogonal components (see Appendix B.7):

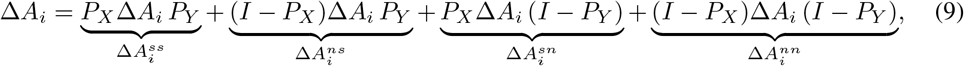

where superscripts denote same (s) or new (n) dimensions relative to the AIS in the source/target respectively, and the score 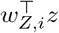 is suppressed (it scales all components equally). Each component has a distinct interpretation (Fig.1c). 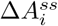: modulation reshapes interactions within the baseline AIS. 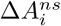: modulation makes source patterns outside the baseline AIS interact with target patterns within it. 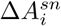: modulation makes target patterns outside the baseline AIS interact with source patterns within it. 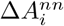: modulation creates new interaction pathways, outside the baseline AIS.

The four components are mutually orthogonal under the Frobenius inner product (Appendix B.7), yielding an additive decomposition of the channel strength 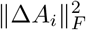. We define fractional *geometry indices* as the fraction of channel strength carried by each component, 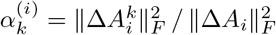 for *k* ∈ {*ss, ns, sn, nn*}, thus having that 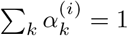. Importantly, the indices depend only on the orientation of *w*_*X,i*_ and *w*_*Y,i*_ relative to the baseline subspaces, not on the modulator, and factorize into independent source and target alignment terms - e.g., 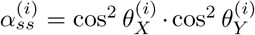, where 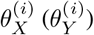 is the angle between *w*_*X,i*_ (*w*_*Y,i*_) and the source (target) AIS - so that all four indices are determined by these two parameters alone (Appendix B.7.2). As a complement, we quantify each component’s contribution to EV_mod_ by fitting constrained MICs that restrict modulation to lie within or outside the baseline subspaces. This reduces to the unconstrained problem on the AIS-projected *X*^*′*^ and *Y* ^*′′*^ (Appendix B.8), enabling direct cross-validated comparison of EV_mod_ across geometric components.

## 4 Validating MICs on simulated data

We validated our pipeline on simulated data (see Appendix B.9 for simulation details).

### 4.1 Rank recovery, sample-efficiency, and benchmarks

To test whether MICs correctly quantify changes in source-target interactions as a function of a modulator *Z*, we first simulated a minimal scenario (Fig. 2a): a 2D source *X* = (*X*_1_, *X*_2_), a 1D target *Y*, and a 1D modulator *Z. X*_1_ drove *Y* additively, independently of *Z* (Fig. 2b), while the *X*_2_ → *Y* interaction was gated by *Z* with strength *γ* (Fig. 2c). AIS correctly recovered a rank-1 subspace from *X* to *Y*, and MICs recovered a rank-1 modulation whenever *γ >* 0 (Fig. 2d). MIC EV increased monotonically with *γ*, while AIS EV decreased as a growing fraction of the variance of *Y* was absorbed by the *Z*-gated interaction - confirming that AIS is blind to modulatory contributions (Fig. 2e). Fitting AIS on *X*_1_ or *X*_2_ alone made this explicit: AIS EV from *X*_1_ matched the full-*X* AIS EV, whereas AIS EV from *X*_2_ remained at zero regardless of *γ* (Fig. 2e, dashed lines).

**Figure 2:**
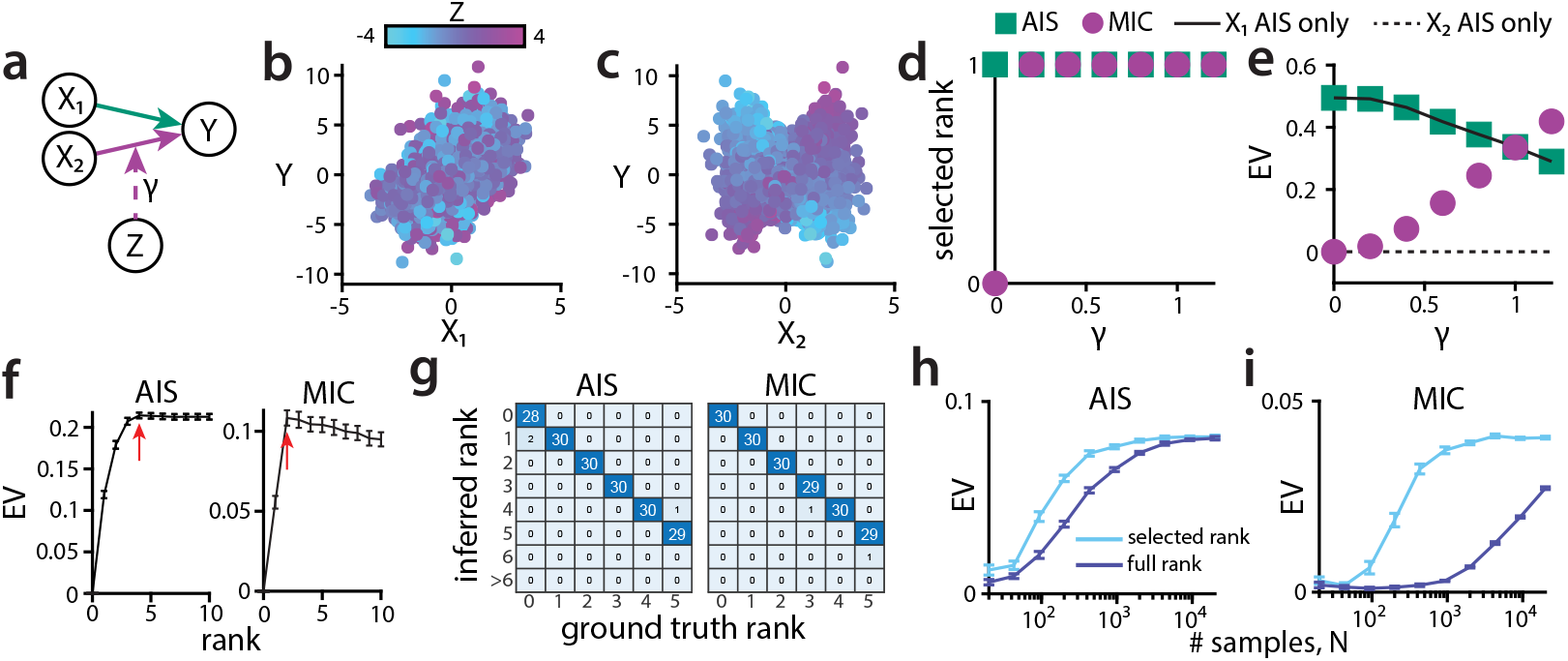
Testing MICs on simulated data. (a) Simulated scenario: a 2D source *X* = (*X*_1_, *X*_2_) interacts with a 1D target *Y* ; the *X*_1_ → *Y* interaction is additive, whereas the *X*_2_ → *Y* interaction is multiplied by *γZ*, where *γ* is the parameter setting the modulation strength. (b) Joint distribution of *X*_1_ and *Y* ; dots color indicates the value of *Z*. (c) Same as (b) for *X*_2_ and *Y* . (d) Selected AIS and modulation ranks as function of *γ*. (e) Explained Variance (EV) by the AIS (green) and MICs (purple) vs *γ*. Solid and dashed black lines denote the AIS EV from *X*_1_ and *X*_2_ alone, respectively. (f) Model rank selection. AIS (left) and modulation (right) EV as function of the fitted rank (ground-truth ranks: *m*_*A*_ = 4, left; *m*_*C*_ = 2, right). Lines: mean ± SEM across cross-validation folds; red arrows: selected rank. (g) Heatmaps of ground-truth versus inferred ranks for the additive (left) and multiplicative (right) channels; *n*_*X*_ = *n*_*Y*_ = *n*_*Z*_ = 20, 5 simulations per (*m*_*A*_,*m*_*C*_) ground-truth rank combination (30 per marginal rank). (h) Variance explained by AIS as a function of sample size *N*, for the rank-selected low-rank model (cyan) and the full-rank model (blue); 30 simulations per scenario, *n*_*X*_ = *n*_*Y*_ = *n*_*Z*_ = 20; *m*_*A*_ = 2, *m*_*C*_ = 1. (i) Same as (h) for MICs.

Next, we considered high-dimensional sources, targets, and modulators (*n*_*X*_ = *n*_*Y*_ = *n*_*Z*_ = 20). We varied the ground-truth AIS and MIC ranks from 0 to 5 drawing *U*_*A*_, *V*_*A*_, *W*_*X*_, *W*_*Y*_, *W*_*Z*_ at random for each simulation. For each fit, we selected the simplest model within 1 SEM of the maximum cross-validated EV (Fig. 2f, [16, 51]). Because this rule occasionally assigns positive ranks when the ground-truth rank is zero (Fig. S1), we retained a non-zero rank only if a residual-shuffle permutation test was significant (*p <* 0.05; Appendix B.13). The full pipeline recovered AIS and MIC ground-truth ranks in most simulations (Fig. 2g). Across all 16 on/off combinations of the four model terms, it yelded 4% and 0% false positives *m*_*A*_ and *m*_*C*_ (Fig. S1a-c). Skipping the significance test inflated them to 18% and 34%, skipping Step 1 inflated *m*_*A*_ whenever *Z* drove both *X* and *Y* (Fig. S1d-i).

We then assessed how AIS and MIC EV recovery scaled with sample size *N* and dimensionality *d* = *n*_*X*_ = *n*_*Y*_ = *n*_*Z*_, against full-rank counterparts (no tensor-rank constraint on AIS or MICs). Reduced-rank fits saturated the ground-truth EV with substantially fewer samples in both cases (Fig. 2h-i and S2a-c, f-h). Normalized recovery curves collapsed onto *N/d* for both reduced-rank models (Fig. S2d, i), versus *N/d*^2^ for full-rank AIS and *N/d*^3^ for full-rank MIC (Fig. S2e, j), consistent with *O*(*d*) per-rank parameter counts against *O* (*d*^2^) and *O* (*d*^3^) for unconstrained AIS and MICs (Appendix B.2). Rank restriction therefore reduces the sample requirement by a factor up to *d* for AIS and *d*^2^ for MIC. Moreover, MICs also tolerated misspecification of the residual noise model: replacing Gaussian emissions on *X* and *Y* with Poisson distribution often used to describe spike counts [52] - or more extreme zero-inflated Poisson preserved EV_mod_ scaling with overall modulation strength and recovery of ground truth MICs direction (Fig. S3).

Since prior work examined how AIS changes across discrete levels of a third variable [3, 19, 53, 54], we benchmarked MICs versus a discretization approach. We partitioned *Z* into *N*_bins_ quantile bins, refitted AIS within each bin, and computed the EV_mod_ as the gap between these binned fits and a single global AIS fit (“Z-split”; same cross-validation folds). For 1D *Z*, EV_mod_ was non-monotonic in *N*_bins_ - rising as finer bins resolved *Z*-dependent structure, then collapsing as per-bin fits became data-starved - but its peak stayed below MIC EV_mod_ (Fig. S4a). For dim(*Z*) *>* 1, where no natural discretization exists, Z-split applied to the leading principal component of *Z* degraded sharply as additional *Z* dimensions misaligned with that component, while MIC EV_mod_ remained stable even in presence of distractor dimensions (Fig. S4b-d).

Finally, we tested MICs on generators that violate bilinearity (Appendix B.10; Fig. S5). In every case EV_mod_ increased monotonically with the modulation strength: a nonlinear ground truth costs MICs sensitivity, not validity. This sensitivity loss is the price of bilinearity. A one-hidden-layer multilayer perceptron fit to the same residuals, with up to 100 times more parameters, captured less modulation than MICs at all sample sizes under bilinear or mildly nonlinear generators, and overtook MICs under strong nonlinearity only at large *N*, at the cost of interpretability (Appendix B.12; Fig. S6).

Together, these results show that MICs efficiently and reliably recover low-dimensional modulation structure in high-dimensional settings - a component of *Y* to which AIS is blind.

### 4.2 Numerical validation of the geometric decomposition

We next tested whether the geometry indices *α* and the constrained-fit EV_mod_ recover the correct MIC geometry with respect to AIS (Fig. 3a). All simulations used *n*_*X*_ = 50, *n*_*Y*_ = 5, and *n*_*Z*_ = 1, implementing a source-target dimensional imbalance which may be present in brain recordings.

**Figure 3:**
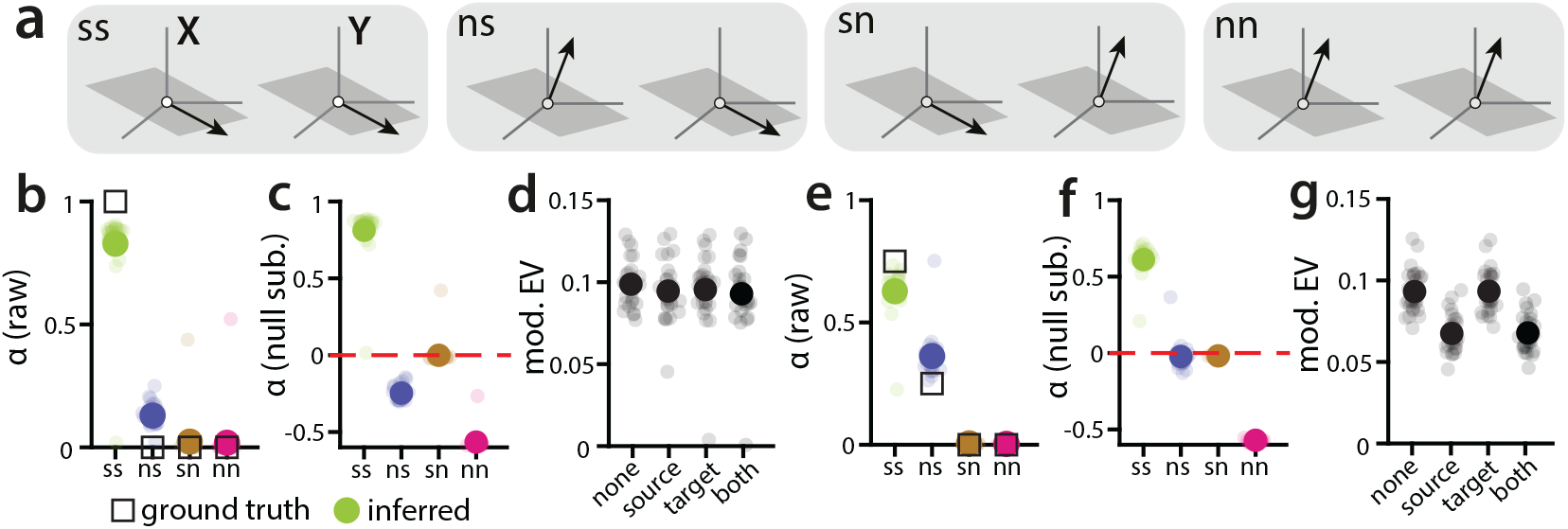
Validating the geometric decomposition on simulations. (a) The four geometric components (*ss, ns, sn, nn*) classifying how *w*_*X,i*_ and *w*_*Y,i*_ align with the baseline AIS (grey plane) on the source and target sides. (b) Raw geometric indices for a scenario with ground-truth (GT) 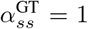 and all others zero. Squares: ground truth; filled circles: mean across simulations; light dots: individual simulations. (c) Same data after subtracting the per-simulation random-rotation null (dashed line: *α* equal to expected value under the null). (d) Constrained-fit EV_mod_ with the indicated MIC weights constrained to lie in the baseline AIS:”source”: *w*_*X*_ ; “target”: *w*_*Y*_ ; “both”: both; “none”: unconstrained fit. (e-g) Same as (b-d) for 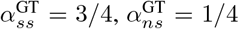. We ran 30 simulations in each scenario with 1000 samples each. *n*_*X*_ = 50, *n*_*Y*_ = 5, *n*_*Z*_ = 1 in all simulations.

We first simulated a MIC entirely confined to the baseline AIS (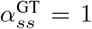, all others zero). Raw indices returned a large *α*_*ss*_ but also a systematic *α*_*ns*_ *>* 0 (Fig. 3b). This positive bias is a geometric artifact of the large difference between *n*_*X*_ and the baseline source rank *m*_*A*_: because *n*_*X*_ − *m*_*A*_ dimensions are available orthogonal to the AIS, small fit-driven deviations of *w*_*X,i*_ out of the baseline subspace inflate *α*_*ns*_. To correct for this, for each fitted MIC we drew *w*_*X,i*_ and *w*_*Y,i*_ independently and uniformly on their respective unit spheres, we recomputed the four indices, and subtracted the random-rotation null means from the raw *α* to obtain null-subtracted 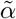 (see Appendix B.7.3). After null subtraction only 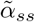 remained positive (Fig. 3c), consistent with ground truth. The constrained-fit EV_mod_ agreed: restricting the MIC to the *ss* component alone (both source and target MICs constrained to the respective AIS) recovered essentially all of the unconstrained EV_mod_ (Fig. 3d). We next considered a mixed geometry (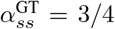 and 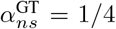). Raw indices correctly individuated both components (Fig. 3e), but null subtraction pushed 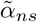 around zero (Fig. 3f): the true *ns* contribution was on the order of the random alignment the null was calibrated to remove, so the test dismissed a genuine component. Constrained EV resolved the ambiguity: the *ss*-only fit recovered ≈ 3*/*4 of the unconstrained EV_mod_. The gap was closed by allowing MICs to lie outside AIS in the source (target constraint; *α*_*ns*_ *>* 0), but not by including *sn* (source constraint; Fig. 3g). A sweep over multi-MIC scenarios (*m*_*C*_ *>* 1) with randomly distributed ground-truth *α* further confirmed that geometry indices recover the correct geometry in more complex settings (Fig. S7). Geometry recovery also held under the nonlinear generators (Fig. S5). Since bilinearity may be violated in *x* or in *z*, we developed a held-out diagnostic that adds higher-order terms in either factor to a fitted channel (Appendix B.11). For sufficient modulation and nonlinearity strength, the diagnostics detected the leading-order nonlinearity of each generator and localized it to the correct factor, and did not detect any nonlinearity on bilinear data (Fig. S8).

Null-subtracted 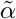 and constrained-fit EV_mod_ are thus complementary: the former is closed-form and cheap, an efficient first-pass diagnostic that can provide false negatives; the latter is more expensive but directly and reliably quantifies each geometric term’s contribution to EV_mod_.

## 5 MICs quantifies stimulus and state modulation of cortical interactions

### Stimulus modulation of V1-V2 interactions (benchmark dataset)

We first applied MICs to the dataset [55] used to benchmark communication subspaces [16]. Anesthetized macaques (3 animals, 5 sessions) were shown drifting gratings with 8 different orientations *θ* while recording activity from V1 and V2 (Fig. 4a; Appendix B.14). Following [16], we analyzed 100 ms spike counts after subtracting the time-dependent trial-averaged activity to eliminate the 1st-order effect of *θ* on both populations (*B* = *D* = 0). Per-orientation communication subspace fits from previous work suggested that the linear map from V1 to V2 residuals depends on *θ*: predictive dimensions fit at one orientation generalize imperfectly to the others [16], and across-area subspaces vary approximately periodically with orientation difference [56]. Both observations rest on comparing separately fitted subspaces, without a statistical test of *θ*-dependence or a measure of its size. We used MICs to fit a single model across all orientations, test this dependence directly, and quantify it. Taking *Z* as a one-hot encoding of *θ*, which imposes no functional form, orientation significantly modulated the interaction in 5/5 sessions (shuffle test, Appendix B.13; EV_mod_ = (1.3 ± 0.1)%, mean ± SEM, i.e. (16.6 ± 2.5)% of the *θ*-invariant EV_AIS_; Fig. 4b, S9a-b; Table 1). Beyond detecting and quantifying the *θ*-dependence of interactions, MICs add two new results. First, it shows the form of the modulation: the modulator scores of an example MIC tracked the angular difference between orientations (Fig. 4c), and the periodic encoding *Z* = (sin 2*θ*, cos 2*θ*) retained (83.6 ± 1.8)% of the one-hot EV_mod_ (Fig. S9d). Thus, the smooth first harmonic accounts for most of a modulation free to take any form, quantifying the periodicity previously observed qualitatively [56]. Second, it determines its geometry relative to the *θ*-invariant AIS, which per-condition refits cannot access: 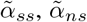 and 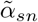 exceeded the null (Fig. 4d, top). The V1- and V2-constrained fits, which add *sn* and *ns* to the both-in-AIS (*ss*-only) fit, respectively, both significantly exceeded it and did not differ significantly from each other, confirming the presence of *ns* and *sn* components (Fig. 4d, bottom). 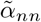 was negative, i.e. the *nn* component was weaker than expected for randomly oriented MICs, but not absent: an *nn*-only model still yielded significant EV_mod_ in 5/5 sessions ((0.24 ± 0.05)% Fig. S9c). The significant *ns* and *sn* components constrain the architecture of V1-V2 interactions. If orientation gated interaction between segregated channels each inside or outside the AIS, modulation would be confined to *ss* and *nn* (Fig. S9f, left). Instead, *θ*-invariant and *θ*-specific interactions are interleaved (Fig. S9f, right): V2 patterns in the AIS interact, at specific orientations, with V1 patterns outside it (*ns*), and vice versa (*sn*).

**Table 1:** Single-session explained variance for the V1-V2 dataset (one-hot encoding of *θ*; all values in %).

| Session | EV<sub>mod</sub> | EV<sub>AIS</sub> | EV<sub>mod</sub>/EV<sub>AIS</sub> |
|---|---|---|---|
| 1 | 1.49 | 10.06 | 14.81 |
| 2 | 1.25 | 9.58 | 13.05 |
| 3 | 1.23 | 10.17 | 12.09 |
| 4 | 0.84 | 5.03 | 16.70 |
| 5 | 1.53 | 5.85 | 26.15 |
| Mean ± SEM | 1.27 ± 0.12 | 8.14 ± 1.11 | 16.56 ± 2.52 |

**Figure 4:**
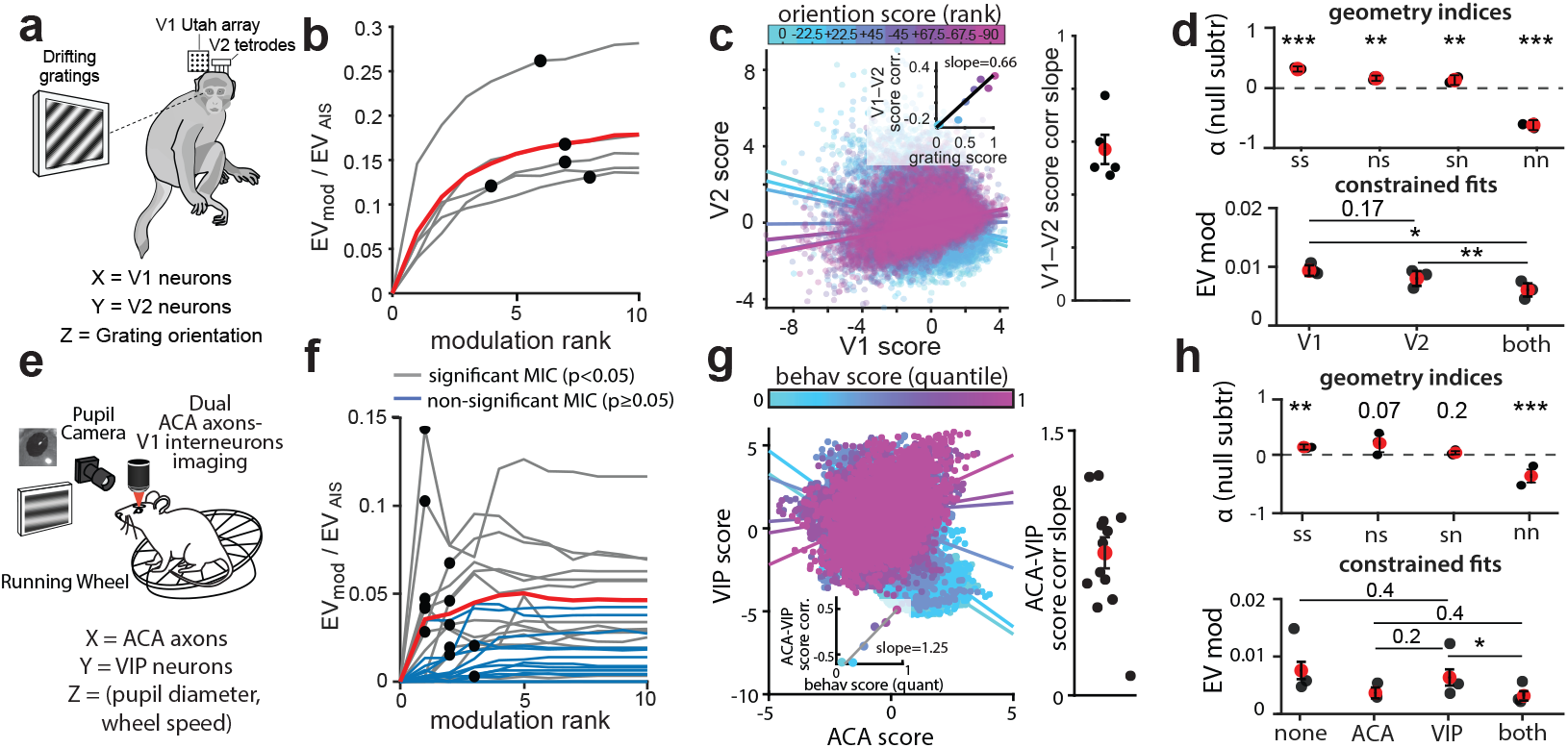
Application of MICs to real data. (a) Simultaneous V1-V2 recordings from macaques shown drifting oriented gratings. (b) Modulatory EV relative to AIS EV vs modulation rank; gray lines: individual sessions (*n* = 5, all with significant EV_mod_); red: mean; black dots: selected rank. (c) Left: V1 vs V2 projection scores of one MIC from an example session. Color: orientation, ranked by its modulator score (one-hot encoding; labels: angular difference from the lowest-scoring orientation). Inset: V1 and V2 scores correlation within each orientation vs orientation score. Right: average correlation slope across MICs; dots: sessions. (d) Top: null-subtracted geometry indices. Bottom: modulatory EV under different geometry constraints (V1: source-in-AIS; V2: target-in-AIS; both: both constraints; n = 3 animals). (e) Simultaneous imaging of ACA axons and VIP neurons in V1 with pupil and wheel measures. (f) Same as (b) for ACA-VIP data; gray (blue) lines: sessions (*n* = 28) with significant (non-significant) EV_mod_. (g) Same as (c) for ACA-VIP data; color: quantiles of behavioral state score. Correlations are computed within each quantile; right: sessions with significant EV_mod_ (*n* = 13). (h) Same as (d) for ACA-VIP data (ACA: source-in-AIS; VIP: target-in-AIS; none: unconstrained; n = 3 animals). P-values in d,h: linear mixed-effects models with animal as random effect (Appendix B.14), testing each index against zero (top), and differences between constrained fits (bottom). *, *p <* 0.05; **, *p <* 0.01; ***, *p <* 0.001. Holm corrected across the comparisons shown, selected as in Appendix B.8. We plot mean ± SEM across sessions.

### Behavioral state modulates prefrontal-visual top-down interactions (new dataset)

Top-down projections from prefrontal cortex (PFC) to V1 gate visual processing in a state-dependent manner [57–59]. Anterior cingulate (ACA, part of PFC) axons modulate responses of Vasoactive Intestinal Peptide (VIP) interneurons in V1 [57], and the activity of both ACA axons and VIP V1 neurons depends on arousal and locomotion [58, 59]. However, whether and how behavioral state reshapes the ACA populations that interact top-down with V1 target populations is unknown. We thus used MICs to address this question. We recorded and publicly release a new dataset of simultaneous two-photon imaging of ACA axonal projections in V1 and V1 VIP interneurons in head-fixed mice during passive visual stimulation (drifting gratings or natural movies), while tracking pupil diameter and running speed (Fig. 4e; Appendix B.15; 3 animals, 28 sessions). We took ACA axons as source *X* and VIP neurons as target *Y* (following anatomical directionality), and defined the modulator *Z* ∈ ℝ ^2^ (hereafter *behavioral state*) as the joint vector of two well-established markers of behavioral state, pupil diameter and running speed [58, 60]. MICs identified significant EV_mod_ in 13/28 sessions (residual circular shuffle test on held-out sequential data halves; Fig. S10a-c; Appendix B.13). In these sessions, MICs accounted for (0.7 ± 0.1)% of total VIP population variance, i.e. (4.7 ± 1.1)% of EV_AIS_ (Table 2; mean selected rank *m*_*C*_ = 1.69). Thus, behavioral state modulates ACA-VIP interactions beyond driving each population alone (Fig. S10d-e), during both grating and natural-movie stimulation (Fig. S10f). The correlation between ACA and VIP MIC projection scores increased with the behavioral-state score 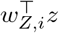 consistently across sessions (Fig. 4g). In contrast to V1-V2, the geometry of these MICs was asymmetric. 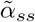 was significantly above zero and 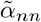 significantly below it, whereas 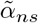 only trended above zero and 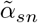 did not (Fig. 4h, top). As the null subtraction can miss a weak genuine component (Fig. 3f), we resolved the geometry with constrained fits (Fig. 4h, bottom). Adding *ns* to the both-in-AIS fit (VIP constraint) significantly increased EV_mod_, whereas adding *sn* (ACA constraint) did not, nor did further adding *sn* and *nn* to the VIP-constrained fit (unconstrained model). Although the ACA- and VIP-constrained fits did not differ significantly, MIC-AIS alignment was significantly larger in VIP than in ACA (Fig. S10g), with MIC weights aligning strongly with the VIP AIS but weakly with the ACA AIS (Fig. S10i-j). Neither this asymmetry nor the geometry indices were explained by the ACA:VIP population-size imbalance (Fig. S10k-m). Both EV_mod_ and the recovered MIC geometry were robust to random initialization (Fig. S11a-b). As in simulations, Z-split fits recovered less than half the EV_mod_ of MICs, degrading further with the number of bins (Fig. S11c-e). Dropping pupil diameter from *Z* did not reduce EV_mod_, whereas dropping running speed did, identifying running speed as the modulating factor (Fig. S11f). Moreover, in both datasets, the bilinearity diagnostic detected no significant nonlinearity in either *x* or *z* (Figs. S9e, S11g). Together, these analyses reveal that behavioral state primarily reconfigures ACA-axon activity patterns interacting with more stable VIP activity patterns.

**Table 2:** Single-session explained variance for the ACA-VIP dataset, for the 13 sessions with significant EV_mod_ (all values in %).

| Session | EV<sub>mod</sub> | EV<sub>AIS</sub> | EV<sub>mod</sub>/EV<sub>AIS</sub> |
|---|---|---|---|
| 1 | 0.82 | 19.02 | 4.31 |
| 2 | 0.27 | 13.20 | 2.05 |
| 3 | 0.07 | 24.10 | 0.29 |
| 4 | 0.57 | 12.13 | 4.70 |
| 5 | 0.79 | 7.65 | 10.33 |
| 6 | 0.99 | 6.88 | 14.39 |
| 7 | 0.16 | 7.98 | 2.01 |
| 8 | 0.19 | 12.24 | 1.55 |
| 9 | 0.42 | 9.21 | 4.56 |
| 10 | 0.67 | 16.06 | 4.17 |
| 11 | 1.49 | 46.01 | 3.24 |
| 12 | 0.89 | 31.35 | 2.84 |
| 13 | 1.95 | 28.88 | 6.75 |
| Mean ± SEM | 0.71 ± 0.15 | 18.05 ± 3.22 | 4.71 ± 1.07 |

## 6 Discussion

MICs extends the CS framework [16] to quantify how a multivariate modulator *Z* reshapes interactions between a source *X* and a target *Y* . Its low-rank structure ensures tractability in the under-sampled, high-dimensional regime of neural data [8, 9]. We provide MICS with a cross-validated hierarchical fitting pipeline a residual-shuffle significance test, a closed-form decomposition into orthogonal geometric components, enabling reliable detection and geometric characterization of whether modulation reshapes baseline interactions, recruits private dimensions of one population, or opens new interactions. This geometric quantification of how a measured modulator changes interactions complements existing models of dynamically modulated population interactions [40–47].

We validated MICs on both simulated and real neural data. On simulated data, MICs reliably recovered ground-truth modulation ranks and MICs geometry with respect to AIS, outperformed simpler approaches, and resolved modulatory EV using up to *d*^2^ fewer samples than full-rank tensor regression. We tested MIC on two neural datasets. On a benchmark existing dataset, MICs recovered the expected stimulus dependence of V1-V2 communication [16, 56] with a single model, resolved its smooth periodic form and the interleaving of stimulus-invariant and stimulus-specific interaction patterns in both areas. On a new dataset that we publicly release (Appendix B.15), MICs revealed that behavioral state reconfigures the interactions between ACA-axon patterns and a stable set of V1 VIP patterns. This asymmetry could reflect a top-down control architecture in which state-dependent recruitment of distinct ACA projection patterns is routed through a conserved VIP interface [57], eliciting a consistent disinhibitory local influence onto V1 pyramidal cells [61].

Several limitations point to natural extensions. MICs capture linear effects of Z on the source-target mapping. The bilinear structure of MICs lends itself to extensions capturing also non-linear effects of Z on the source-target mapping [47], and to learning the modulator encoding function from data. Moreover, so far we restricted our analyses to interactions at zero time lag, with source and target assigned from anatomy. Similar to time-lagged CS analyses [56], using MICs with time-lagged-interactions, possibly combined with Wiener-Granger analyses [11, 12], would allow MICs to assess from data whether interactions are unidirectional or bidirectional, and to quantify how each direction is modulated. In addition, our significance test fits on one half of a session and tests generalization to the other, entirely held-out half. This makes it conservative and data-demanding. Thus more data-efficient tests will need to be developed. Finally, conclusions from our real datasets are restricted to the small populations of recorded animals. Further, we currently take the baseline as the session mean. Alternative schemes (e.g., anchoring the baseline at biologically interpretable values of the modulator) are possible. Finally, the subspaces jointly estimated by MICs for encoding (*B, D*) and interaction (AIS), possibly integrated with other tools to study the geometry of population coding ([62–64]), could address whether the population activity subspaces encoding information coincide or differ from those that transmit information about the modulating variable across areas, providing a unifying framework for information encoding and transmission.

By providing a compact, data-efficient, and theoretically grounded characterization of how a possibly high-dimensional modulator reshapes interactions between high-dimensional populations, MICs are well positioned to enable new discoveries about context-dependent interactions in neural systems.

## Acknowledgments and Disclosure of Funding

We thank S.M. Lemke and members of the Sur lab for feedback and helpful discussions, and G. Sipe for the illustration in Fig. 4e. This work was supported by European Union’s Horizon research and innovation programme grant agreement No. 101206609, Marie Sklodowska-Curie Global Fellowship - AstroError (MC); the German Federal Ministry of Education and Research (BMBF 01GQ2404 and 01DM26028; SP); NIH grants R01MH126351; R01MH133066; R01NS130361; R01MH085802; MURI Grant W911NF2110328; The Picower Institute Innovation Fund; and the Simons Foundation Autism Research Initiative through the Simons Center for the Social Brain (MS); NIH grant 1K99EY035752 (SÄR); NIH grant F32EY032756 (KRJ); the MIT MSRP Bio program (NSF STC award CCF-1231216; JSS).

## A Glossary

In this paper, we use the following abbreviations:

| Abbreviation | Meaning |
|---|---|
| RRR | Reduced-rank regression. |
| CS | Communication subspace: the framework of [16], which uses RRR to identify covarying low-dimensional activity patterns of a source and a target population. |
| AIS | Additive interaction subspaces: the source and target subspaces estimated within the CS framework; renamed to emphasize that they capture additive covariation, not directed communication. |
| MIC | Multiplicative interaction channel: a triplet of source, target, and modulator patterns, in which the modulator pattern multiplicatively scales the interaction between the source and target patterns. |
| MICs | Our method, extending the CS framework to infer, within a single model, the subspaces encoding the modulator, the additive interaction subspaces, and the multiplicative interaction channels. |
| CP | Canonical Polyadic (tensor decomposition into a sum of rank-1 terms). |
| ALS | Alternating least squares. |
| EV | Explained variance. |
| GT | Ground truth. |
| IQR | Interquartile range. |
| MLP | Multilayer perceptron. |
| ReLU | Rectified linear unit. |
| SEM | Standard error of the mean. |
| ACA | Anterior cingulate area. |
| PFC | Prefrontal cortex. |
| V1, V2 | Primary and secondary visual cortex. |
| VIP | Vasoactive intestinal peptide-expressing interneurons. |
| PSTH | Peri-stimulus time histogram (time-dependent firing rate averaged over all trials for a given stimulus). |
| ROI | Region of interest. |

## B Technical appendices

Throughout the appendix, we use the notation established in the main text: 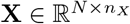(source), 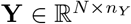 (target), 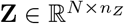 (modulator), all zero-centered column-wise. We use ∥ · ∥_*F*_ for the Frobenius norm, ⟨*K, L*⟩_*F*_ = tr(*K*^*⊤*^*L*) for the Frobenius inner product, ⊙ for the element-wise (Hadamard) product, and ⊗ for the outer (tensor) product.

### B.1 Introduction to reduced-rank regression

We briefly summarize the ridge-regularized reduced-rank regression (ridge-RRR) estimator used throughout this work. For full derivations, see [48, 17].

Consider the linear model **Y** = **X***A* + **E**, where 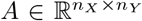 with rank(*A*) ≤ *m*. We seek to minimize the ridge-regularized loss subject to a rank constraint:

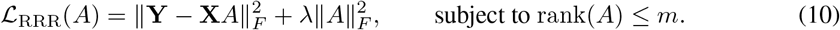

#### Full-rank ridge estimate

Ignoring the rank constraint, setting ∇_*A*_ ℒ = 0 yields:

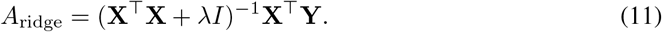

#### Rank-constrained projection

The key result (see [17], Appendices C-D, for a complete proof) is that incorporating the rank constraint is well approximated by projecting the ridge estimate onto the subspace spanned by the top *m* eigenvectors of a specific matrix. Concretely, the ridge-RRR solution is:

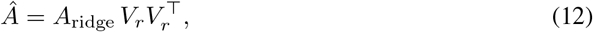

where 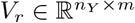 contains the top *m* eigenvectors of the symmetric positive semidefinite matrix 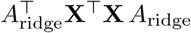. Equivalently, *V*_*r*_ is obtained by performing PCA on the *N* × *n*_*Y*_ matrix of ridge-regularized predictions **X** *A*_ridge_ and retaining the top *m* components.

#### Interpretation: source and target subspaces

The solution in Eq. 12 has the factorized form 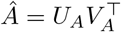, with:

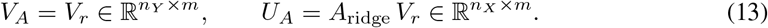

These factors have direct interpretations in terms of the relationship between source and target variables:

- **Source AIS**, 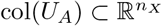 : the *m*-dimensional subspace of source patterns that co-vary with the target. Source patterns orthogonal to this subspace are “private” - they have no linear relationship with the target.
- **Target AIS**, 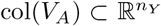: the *m*-dimensional subspace of target patterns that can be linearly related to the source. Target patterns orthogonal to this subspace are “private” and unexplained by the source.

Note that while the subspaces col(*U*_*A*_) and col(*V*_*A*_) are uniquely determined by *Â*, the individual columns of *U*_*A*_ and *V*_*A*_ are not: for any invertible *L* ∈ ℝ^*m×m*^, the alternative factorization *Â* = (*U*_*A*_*L*)(*V*_*A*_*L*^*−⊤*^)^*⊤*^ yields the same product. This rotational ambiguity means that individual “communication dimensions” within the subspace are not identifiable from the model alone.

### B.2 From scalar interaction regression to the MICs modulation tensor

In this section, we position the MICs model within the broader landscape of interaction modeling: we start from scalar regression with an interaction term, generalize to the high-dimensional setting where the interaction is encoded by a three-way tensor *C*, recover MICs as a CP-rank constraint on *C*, and finally relate it to prior tensor regression. To build intuition, we briefly connect our model to the familiar setting of regression with interaction terms. Consider first a scalar source *x* ∈ ℝ, scalar modulator *z* ∈ ℝ, and scalar target *y* ∈ ℝ. A standard interaction model is:

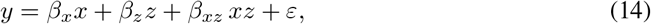

where *β*_*xz*_ *xz* is the interaction term: the effect of *x* on *y* depends linearly on *z*, and vice versa.

Our model generalizes this to the high-dimensional setting where 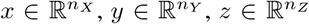 . Without any dimensionality reduction, the target for observation *n* would be:

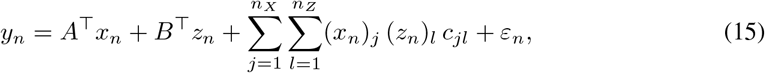

where 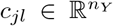 is a vector of interaction coefficients for source dimension *j* and modulator dimension *l*. This full interaction model has *n*_*X*_ × *n*_*Z*_ × *n*_*Y*_ free parameters in the interaction term alone, which is prohibitively large in typical neuroscience settings (e.g., *n*_*X*_ = *n*_*Y*_ = *n*_*Z*_ = 50 gives 125,000 interaction parameters).

Equivalently, collecting the vectors 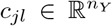into a single three-way array, i.e. *the full-rank modulation tensor*, 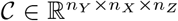 with entries *C*_*i,j,l*_ = (*c*_*jl*_)_*i*_, the modulation term in Eq. 15 is the contraction of *C* against the source and modulator vectors of observation *n*:

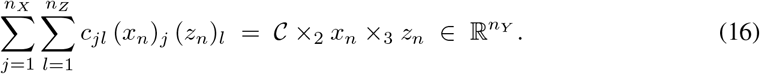

Here ×_*k*_ denotes the *mode-k vector product*, i.e., contraction of a tensor along its *k*-th index: for 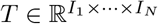 and 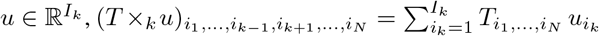 [33]. Mode 2 (size *n*_*X*_) is thus contracted with the source *x*_*n*_ and mode 3 (size *n*_*z*_) with the modulator *z*_*n*_, leaving mode 1 (size *n*_*Y*_) as the output. Equivalently, *C* is evaluated on the rank-1 predictor 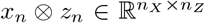 formed by source and modulator at observation *n*.

The MICs model (Eq. 5) greatly reduces the number of estimated parameters by imposing a low-rank structure on C. Each MIC *i* contributes a rank-1 interaction:

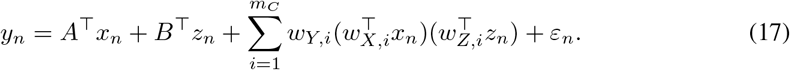

This is exactly Eq. 15 with the three-way array of interaction coefficients constrained to have the CP decomposition 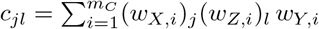, reducing the number of interaction parameters from *n*_*X*_ × *n*_*Z*_ × *n*_*Y*_ to *m*_*C*_(*n*_*X*_ + *n*_*Z*_ + *n*_*Y*_). For *m*_*C*_ = 3 and *n*_*X*_ = *n*_*Y*_ = *n*_*Z*_ = 50, this is a compression from 125,000 to 450 parameters. Importantly, the low-rank structure is not merely a computational convenience: it provides interpretability. Each channel defines a specific source axis (*w*_*X,i*_), modulator axis (*w*_*Z,i*_), and target axis (*w*_*Y,i*_), making it possible to ask which specific source patterns have their relationship with which specific target patterns modulated by which specific modulator patterns. In the scalar case (Eq. 14), there is only one such “channel” (*m*_*C*_ = 1, with *w*_*X*_ = *w*_*Z*_ = *w*_*Y*_ = 1, and the channel strength equals *β*_*xz*_).

In isolation, the modulation term is a specific instance of the tensor-on-tensor regression framework of [38], with per-observation predictor tensor 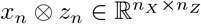 and outcome 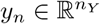 . The rank-1 structure of the predictor - the outer product of two distinct observed variables rather than a single multiway array - is what endows the CP decomposition with its bilinear-interaction interpretation. The full MICs model (Eq. 5) embeds this term inside an additive RRR framework with residualized source *X*^*′*^ = *X* − *ZD*, which falls outside [38]: Lock’s framework fits a single coefficient tensor governing one predictor, and does not address the disentangling of additive from multiplicative effects of distinct predictors *X* and *Z*. Beyond the model itself, the hierarchical pipeline (Algorithm 1) handles the simultaneous presence of additive and multiplicative effects of Z under shared cross-validation folds, and the geometric decomposition (Section 3.4) exploits the existence of a baseline AIS operator A against which each MIC is projected - neither of which has a counterpart in [38].

### B.3 Uniqueness of MICs: Kruskal’s theorem

The use of *channel* rather than *subspace* to describe MICs reflects an algebraic distinction from AIS. Reduced-rank matrix factorizations identify column subspaces but not individual factor columns, which carry rotational indeterminacy. The three-way tensor structure of *C* breaks this indeterminacy: under a condition on the Kruskal ranks of the factor matrices, the individual rank-1 components of the CP decomposition are essentially unique, and identifiable up to permutation and per-channel rescaling (Kruskal’s theorem). We state the theorem, verify the condition in our setting, and note its practical consequences for MIC analysis.

#### B.3.1 Kruskal rank and theorem statement

##### Definition B.1.

The *Kruskal rank* of a matrix *M* ∈ ℝ^*n×r*^, denoted *k*(*M*), is the largest integer *j* such that every subset of *j* columns of *M* is linearly independent.

Note that *k*(*M*) ≤ rank(*M*) ≤ min(*n, r*), with equality *k*(*M*) = rank(*M*) when every subset of rank(*M*) columns is linearly independent. For a matrix whose columns are drawn independently from a continuous distribution on ℝ^*n*^ with *n* ≥ *r, k*(*M*) = *r* with probability one.

##### Theorem B.2

(Kruskal, 1977 [49]). *Let* 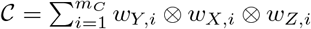 *be a rank-m*_*C*_ *CP decomposition with factor matrices* 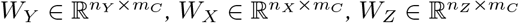 . *If*

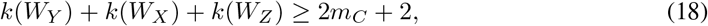

*then the decomposition is* essentially unique: *any other rank-m*_*C*_ *decomposition is related to it by a permutation of the channels and per-channel rescaling*.

In settings where, *n*_*X*_, *n*_*Y*_, *n*_*Z*_ ≫ *m*_*C*_ and the weight vectors are estimated from data with continuous noise, generically *k*(*W*_*Y*_) = *k*(*W*_*X*_) = *k*(*W*_*Z*_) = *m*_*C*_. Eq. 18 then reduces to 3*m*_*C*_ ≥ 2*m*_*C*_ + 2, i.e., *m*_*C*_ ≥ 2 (uniqueness for *m*_*C*_ = 1 is trivial up to an overall scalar). This argument breaks when any factor dimension is small relative to *m*_*C*_. For example, *k*(*W*_*Z*_) = 1 for a scalar modulator (*n*_*Z*_ = 1), so Eq. 18 fails for all *m*_*C*_ ≥ 2.

#### B.3.2 Practical implications for MIC analysis

Many natural analyses with AIS are subspace-level - for example, predicting the target response evoked by a given source pattern - and do not require identifiability of individual factor columns. Kruskal’s theorem provides a stronger guarantee that becomes relevant when multiple MICs with distinct geometries coexist in the same population. Without column-level identifiability, two channels with different geometric profiles (e.g., one predominantly *α*_*ss*_, another predominantly *α*_*ns*_) could be rotated into mixtures, blurring their distinct alignment with the baseline AIS. Theorem B.2 ensures that this cannot happen: each channel retains its individual geometry. We do not fully exploit this property in the present manuscript - our current analyses summarize geometric structure at the level of MIC-averaged indices - but it opens the door to per-channel comparative analyses (e.g., contrasting the biological correlates of channels with different *α* profiles) that we view as a promising direction for future work.

The CP parametrization has an inherent per-channel scaling ambiguity: (*w*_*X,i*_, *w*_*Z,i*_, *w*_*Y,i*_) and (*w*_*X,i*_*/β, βγ w*_*Z,i*_, *w*_*Y,i*_*/γ*) define the same channel, since the outer product is multilinear. Under the identifiability conditions of Theorem B.2, this is the *only* remaining ambiguity. We resolve this by extracting unit-norm directions and absorbing the magnitude into a single *channel strength σ*_*i*_ = ∥*w*_*X,i*_∥ · ∥*w*_*Z,i*_∥ · ∥*w*_*Y,i*_∥, giving 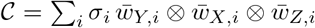. This normalization is used throughout the paper when displaying or comparing MIC weights and scores (e.g., Fig. 4c, Fig. S10j-k). It does not affect the geometry indices *α* (which depend only on directions) or the channel’s contribution to EV_mod_.

### B.4 The CP Frobenius penalty

The modulation tensor 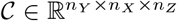, with CP decomposition 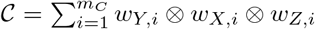 has squared Frobenius norm:

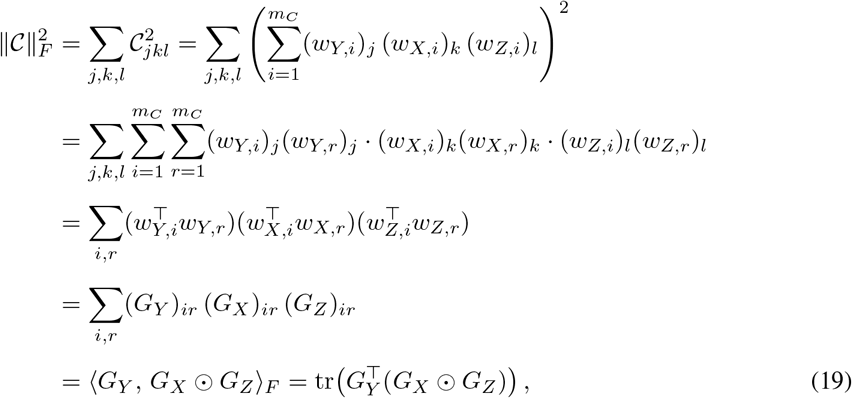

where 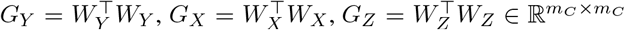 are the Gram matrices of the factor matrices, and the last line uses the identity ∑_*ir*_ *A*_*ir*_*B*_*ir*_*C*_*ir*_ = ⟨*A, B* ⊙ *C*⟩_*F*_ for the element-wise product. This expression is efficient to compute (it involves only *m*_*C*_ *× m*_*C*_ matrices) and is used in the regularization term of the modulation loss (Eq. 8 in the main text).

### B.5 ALS derivation for the modulation term

In this section, we derive ALS for the modulation loss of our model (Eq. 8). ALS for ridge-regularized CP-rank tensor regression is an established tool [33, 37, 38]. The closed-form updates derived below follow from specializing this framework - specifically the non-separable *L*_2_ penalty 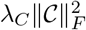on the CP coefficient tensor of [38] - to the bilinear structure of Eq. 8. We provide the derivation for self-containedness and summarize it in algorithm 2.

The modulation loss of our model with the penalty term expressed as in Eq. B.4 is:

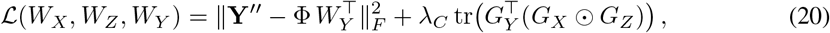

where 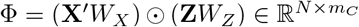, and **Y**^*′′*^ = **Y**^*′*^− **X**^*′*^*A* are the residuals from the additive model. Each ALS sweep updates *W*_*Y*_, then *W*_*X*_, then *W*_*Z*_, while holding the other two fixed, similarly to the ALS procedures for CP-rank-constrained tensor regression developed in [37, 38].

Expanding the first terms in the RHS of Eq.20 yields:

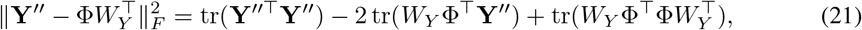

and, in the following, we refer to the two terms that depend on the factor matrices as the cross-term −2 tr(*W*_*Y*_ Φ^*⊤*^**Y**^*′′*^) and the quadratic term 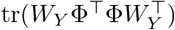.

#### B.5.1 Update for *W*_*Y*_

With *W*_*X*_ and *W*_*Z*_ (and hence Φ, *G*_*X*_, *G*_*Z*_) fixed, the loss is quadratic in *W*_*Y*_ . Taking the derivative of Eq.20 with respect to 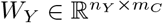 and setting it to zero yields:

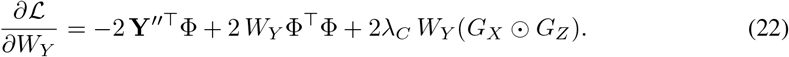

The derivative of the penalty term (i.e. the second term in the RHS of Eq.20) follows from 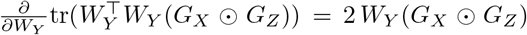, since *G*_*X*_ ⊙ *G*_*Z*_ is symmetric. Setting the 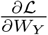 derivative to zero gives:

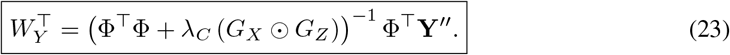

This is an *m*_*C*_ × *m*_*C*_ linear system (one right-hand side per target dimension), which is inexpensive to solve.

#### B.5.2 Update for *W*_*X*_

With *W*_*Z*_ and *W*_*Y*_ fixed, the columns of *W*_*X*_ are coupled through both the Hadamard product in Φ and the CP penalty. We derive the gradient via total differentials.

Define 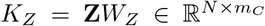 and 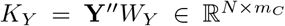. Considering a perturbation *W*_*X*_ → *W*_*X*_ + *dW*_*X*_, we compute the three contributions to *dℒ* coming from the cross-term, the quadratic term and the penalty term respectively:

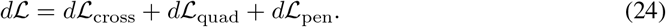

i. *Cross term* −2 tr(*W*_*Y*_ Φ^*⊤*^**Y**^*′′*^): Under *d*Φ = (**X**^*′*^ *dW*_*X*_) ⊙ *K*_*Z*_ and using the Hadamard product identity ⟨*A, B* ⊙ *C*⟩_*F*_ = ⟨*A* ⊙ *B, C*⟩_*F*_ :

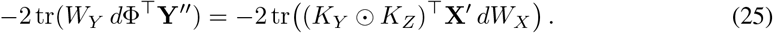
ii. *Quadratic term* tr(*G*_*Y*_ Φ^*⊤*^Φ):

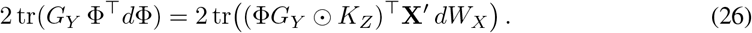
iii. *Penalty term:* Since 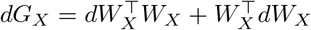,

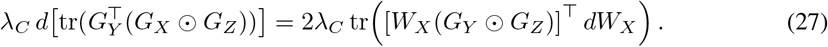

Therefore,

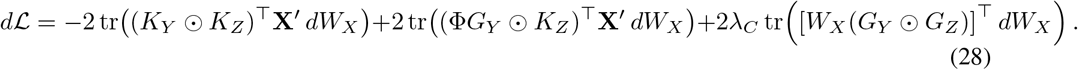

Combining and identifying the gradient of the scalar loss via 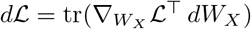:

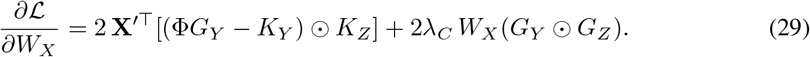

##### Block linear system

Setting Eq. 29 to zero does not yield a decoupled update for the columns of *W*_*X*_, because the columns are coupled through Φ*G*_*Y*_ and through the penalty term. Let *L* = *G*_*Y*_ ⊙*G*_*Z*_ and 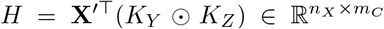. Writing 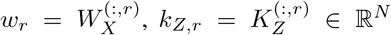, and *h*_*j*_ = *H*^(:,*j*)^, the stationarity condition for the *j*-th column is

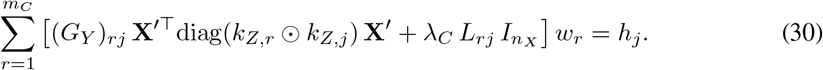

where 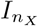 denotes the *n*_*X*_ *× n*_*X*_ identity matrix. Stacking the columns of *W*_*X*_ yields the linear system

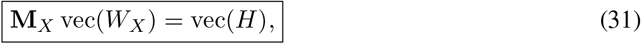

where 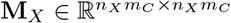 is a symmetric block matrix with (*j, r*)-th block (*n*_*X*_ × *n*_*X*_ each)

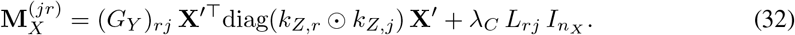

The diagonal matrix need not be formed explicitly: for any *v* ∈ R^*N*^,

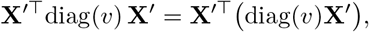

i.e. by scaling the rows of **X**^*′*^ by *v* before left-multiplying by **X**^*′⊤*^.

#### B.5.3 Update for *W*_*Z*_

By symmetry between **X**^*′*^ and **Z** in the Hadamard product Φ = (**X**^*′*^*W*_*X*_) ⊙ (**Z***W*_*Z*_), the update for *W*_*Z*_ is obtained by swapping (**X**^*′*^, *W*_*X*_, *G*_*X*_) with (**Z**, *W*_*Z*_, *G*_*Z*_). Defining *K*_*X*_ = **X**^*′*^*W*_*X*_ (using the just-updated *W*_*X*_), *L*^*′*^ = *G*_*Y*_ ⊙ *G*_*X*_, and *H*^*′*^ = **Z**^*⊤*^(*K*_*Y*_ ⊙ *K*_*X*_):

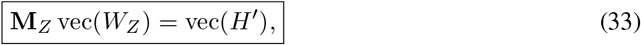

with (*j, r*)-th block

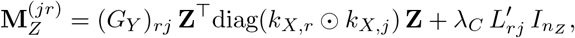

where 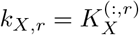.

It is worth noticing that the ALS objective in Eq. 20 is non-convex. In principle, different initializations may converge to different local optima. Empirically, on the ACA-VIP data both EV_mod_ and the recovered MIC geometry were stable across random ALS initializations (Fig. S11; Appendix B.15).

### B.6 Hierarchical model fitting algorithm

For clarity, we summarize the full fitting pipeline in Algorithm 1. The algorithm mirrors the structure of the implementation described in the main text and in the code: first fit the additive effects of the modulator on source and target, then fit the baseline AIS on the residualized variables, and finally fit the MICs on the residuals of the baseline model.

In the pseudocode below, calligraphic symbols denote collections of fold-specific quantities across cross-validation splits. For example, 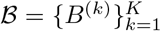 denotes the collection of *Z* → *Y* regression matrices obtained from each fold *k*, and 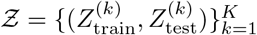 denotes the collection of train/test splits of *Z*. Algebraic expressions involving calligraphic variables indicate fold-wise operations. For instance,

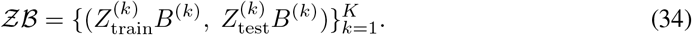

Algorithm 1 gives the global structure of the fitting pipeline. The final step, CFit, requires solving the low-rank tensor regression problem for the modulation term. We summarize the complete ALS procedure used inside CFit in Algorithm 2. The weight matrices are initialized randomly (from a standard normal distribution), and sweeps are repeated until the relative change in the loss falls below a tolerance *ϵ* or a maximum number of sweeps *maxIter* is reached (in our implementation we set *ϵ* = 10^*−*5^ and *maxIter* = 100).

The computational cost per sweep is dominated by assembling and solving the *n*_*X*_ *m*_*C*_ × *n*_*X*_ *m*_*C*_ and *n*_*Z*_*m*_*C*_ × *n*_*Z*_*m*_*C*_ block systems for *W*_*X*_ and *W*_*Z*_: *O*((*n*_*X*_ *m*_*C*_)^2^*N* + (*n*_*X*_ *m*_*C*_)^3^) and analogously for *W*_*Z*_. The *W*_*Y*_ update costs only 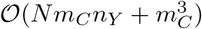

#### Algorithm 1

Hierarchical fitting procedure for MICs

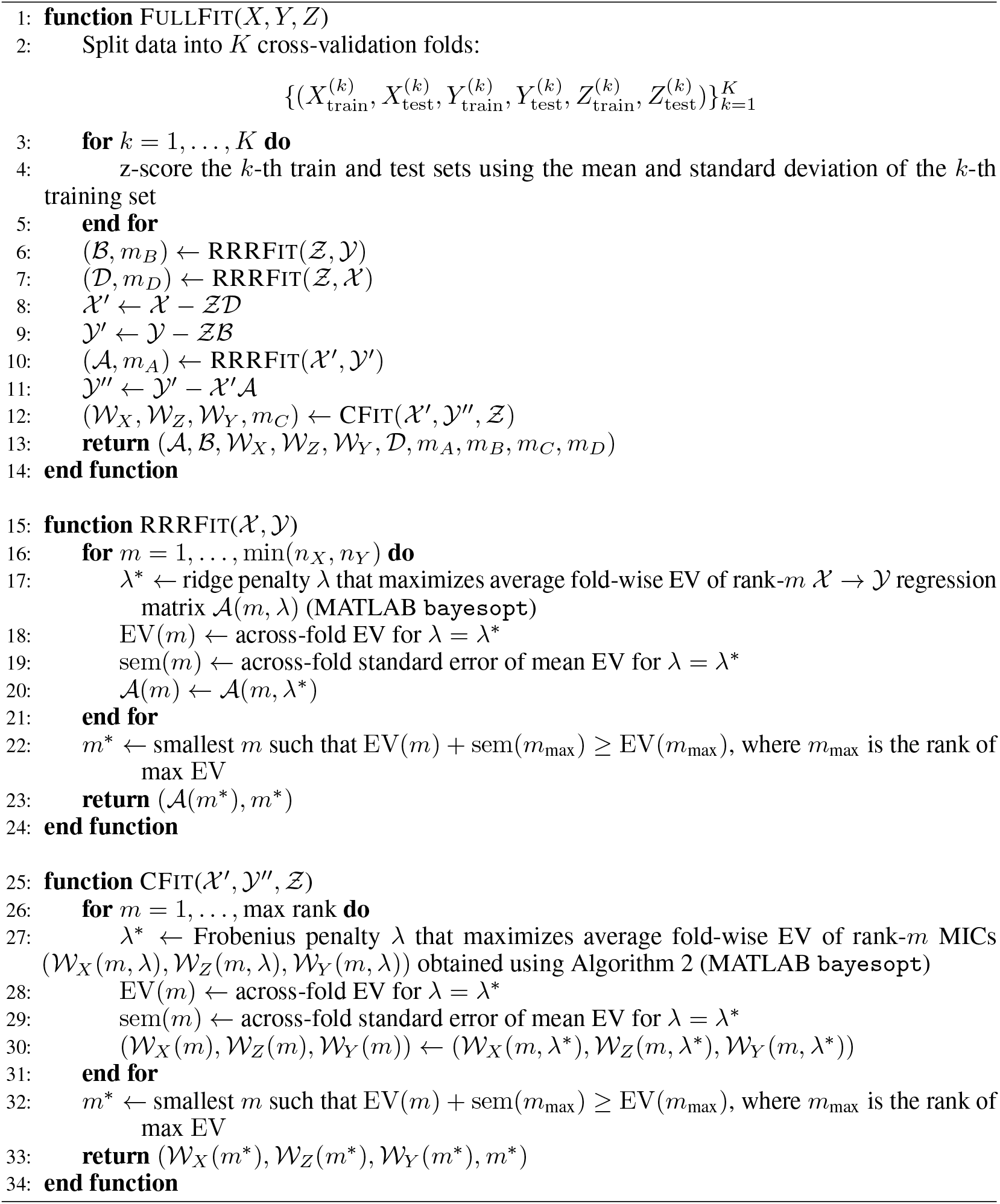

#### Modulatory explained variance

Denoting by 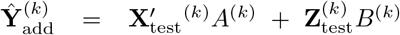 the joint additive model prediction on the test set of fold *k*, and by 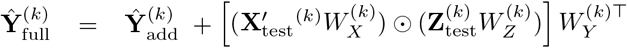 the full model prediction, the modulatory explained variance (Sec. 3.3) is

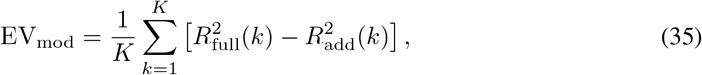

where *R*^2^(*k*) is the coefficient of determination on the test set of fold *k*. Analogously, EV_AIS_ is the fold-averaged *R*^2^ increment of 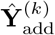 over the modulator-only prediction 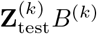.

##### Algorithm 2

ALS for fitting MICs

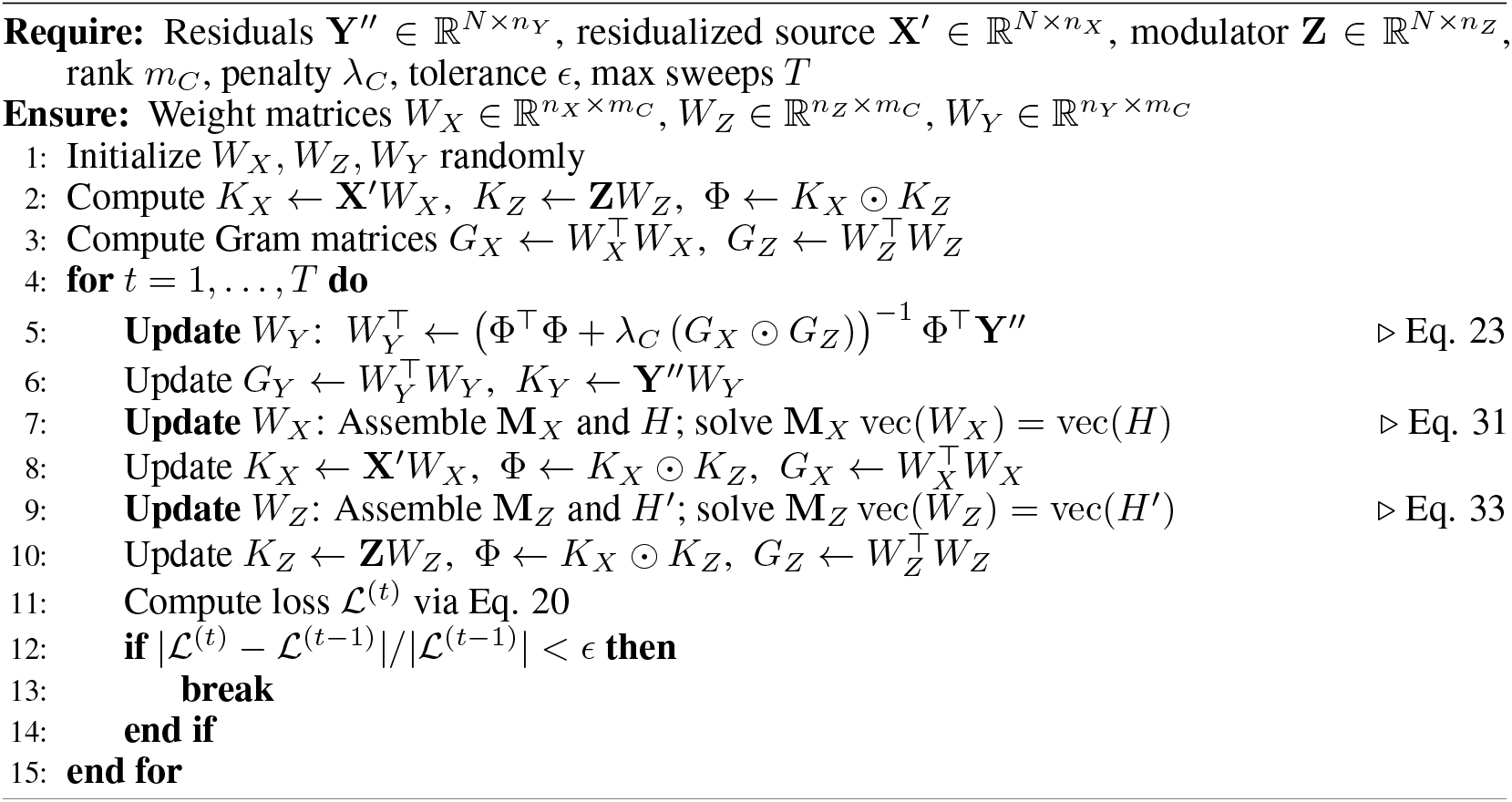

### B.7 Geometric decomposition of MICs

In this section, we derive the four-component decomposition of each channel Δ*A*_*i*_ and the resulting form of the geometry indices:

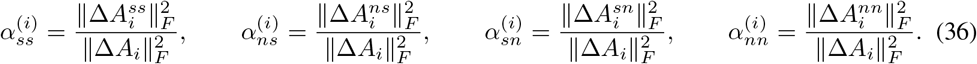

The results in this section follow from standard algebra of orthogonal projectors (see, e.g., [65]) and are reported here for clarity and completeness; we do not claim them as novel theoretical contributions.

#### B.7.1 Orthogonal decomposition

The decomposition *M* = *M* ^*ss*^ + *M* ^*ns*^ + *M* ^*sn*^ + *M* ^*nn*^ (Eq. 9) follows by inserting *I* = *P* + (*I* − *P*) on both sides of *M* :

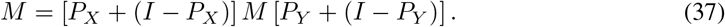

The pairwise orthogonality of the four components under the Frobenius inner product is a consequence of the idempotence of orthogonal projectors. For any two components that differ in their source-side projector (e.g., *M* ^*ss*^ and *M* ^*ns*^):

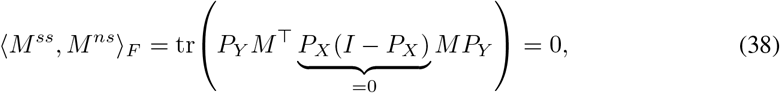

and for components that differ in their target-side projector (e.g., *M* ^*ss*^ and *M* ^*sn*^), the analogous identity using (*I* − *P*_*Y*_)*P*_*Y*_ = 0 and the cyclic property of the trace applies. The Pythagorean norm decomposition 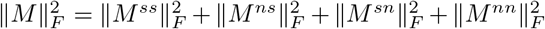 is an immediate consequence.

#### B.7.2 Explicit form of *α* indices for rank-1 perturbations

For a rank-1 matrix 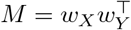, the four components remain rank-1 (or zero):

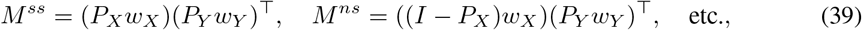

since orthogonal projectors act linearly: 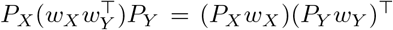. Using the identity 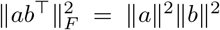 for rank-1 matrices (which follows from 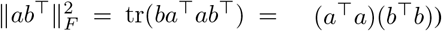, the squared Frobenius norms factorize:

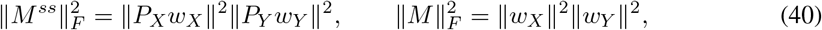

and analogously for the other components. The geometry indices (Eq. 36) therefore take the factorized form:

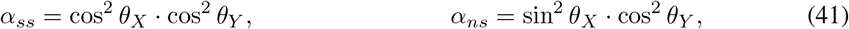

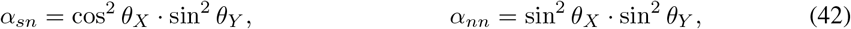

where cos^2^ *θ*_*X*_ = ∥*P*_*X*_ *w*_*X*_ ∥^2^*/* ∥*w*_*X*_ ∥^2^ is the squared cosine of the angle between *w*_*X*_ and the source AIS, and cos^2^ *θ*_*Y*_ = ∥*P*_*Y*_ *w*_*Y*_ ∥^2^*/* ∥*w*_*Y*_ ∥^2^ analogously for the target. The Pythagorean identity for projections ensures cos^2^ *θ* + sin^2^ *θ* = 1 on each side, so the four *α* values sum to one.

##### Remark B.3

(Independence from the modulator). Since each MIC *i* contributes 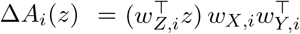, the scalar 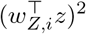 appears in both numerator and denominator of each *α* index and cancels. The geometry indices thus characterize a fixed geometric property of each channel, regardless of the instantaneous value of *z* or the modulator weight vector *w*_*Z,i*_.

##### Remark B.4

(Two-parameter structure). The product form reveals that the full 2 × 2 table of *α* values is determined by just two free parameters: the source alignment cos^2^ *θ*_*X*_ and the target alignment cos^2^ *θ*_*Y*_ . Defining *a*_*X*_ = cos^2^ *θ*_*X*_ and *a*_*Y*_ = cos^2^ *θ*_*Y*_ :

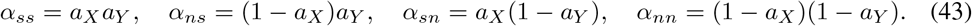

#### B.7.3 Null randomized test of *α* indices

Raw geometry indices *α*_*k*_ are subject to a systematic bias: finite-sample estimation causes the fitted directions **w**_*X,i*_, **w**_*Y,i*_ to deviate from their true orientations, and the dimensional structure of the problem (*n*_*X*_ ≫ *m*_*A*_ in typical recordings) determines where those deviations land - predominantly outside the AIS, inflating components such as *α*_*ns*_ and *α*_*nn*_ even when the true geometry is confined to *α*_*ss*_ (Fig. 3b).

To correct for this, we estimate the expected *α* values under a *random-direction null*: for each fitted MIC *i*, we replace **w**_*X,i*_ and **w**_*Y,i*_ with independent draws from the uniform distribution on their respective unit spheres, recompute the four indices, and repeat for *N*_shuf_ shuffles. The null-subtracted index is

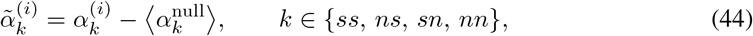

where ⟨·⟩ denotes the mean over shuffles.

Values 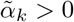 indicates that the fitted MIC aligns with geometric component *k* more than expected for a uniformly random direction in the same ambient space - that is, the channel exhibits geometric structure in that component beyond chance. The test is conservative: components whose true weight is small relative to the random-alignment baseline will be dismissed Fig. 3f), in which case constrained-fit EV_mod_ Appendix B.8) provides a more sensitive characterization of MICs geometry.

### B.8 Constrained fitting: modulation within or outside baseline subspaces

As described in Section 4.2, we test whether modulatory variance is attributable to specific geometric components by fitting constrained models. The following result shows that constraining MIC factors to specified subspaces reduces, up to a constant, to an unconstrained problem on projected data, so that Algorithm 2 can be reused without modification.

#### B.8.1 General reduction

##### Proposition B.5

(Constrained fitting reduction). *Let* 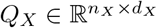 *and* 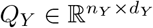 *be semi-orthogonal matrices* 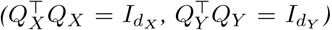, *and consider the modulation loss (Eq. 20) constrained by the parametrization*

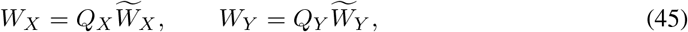

*with* 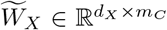 *and* 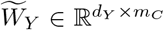 *free. Then the constrained loss is equivalent, up to an additive constant independent of the free parameters, to the unconstrained modulation loss applied to the projected data* (**X**^*′*^*Q*_*X*_, **Y**^*′′*^*Q*_*Y*_, **Z**).

*Proof*. Substituting Eq. 45 into the modulation loss and splitting the data-fit term using the fact that the prediction **Ŷ** lies entirely in col(*Q*_*Y*_):

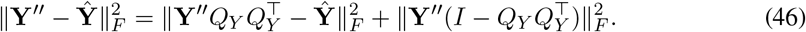

The second term is constant in the free parameters. For the first, 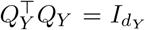 gives 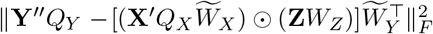 . The penalty simplifies analogously: 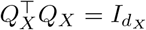 and 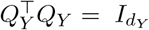 give Gram matrices 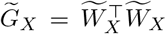 and 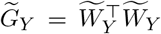, so 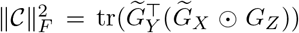. Combining:

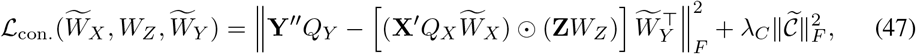

which is the unconstrained modulation loss in the projected variables. □ □

Eq. 47 can be minimized using the same ALS procedure (Algorithm 2). After optimization, the full-space factors are recovered via 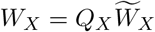 and 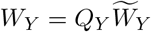.

#### B.8.2 Specific geometric constraints

The geometric components of the *α* decomposition correspond to specific choices of (*Q*_*X*_, *Q*_*Y*_). We list all eight possible constraints below for completeness. In the present paper we use four of these constraints, compared with each other and with the unconstrained fit (Sections 3.4 and 5): the both-in-AIS (*α*_*ss*_-only) fit, the source-in-AIS fit (*w*_*X*_ in the source AIS, *w*_*Y*_ free; forces *α*_*ns*_ = *α*_*nn*_ = 0), the target-in-AIS fit (*w*_*Y*_ in the target AIS, *w*_*X*_ free; forces *α*_*sn*_ = *α*_*nn*_ = 0), and the *α*_*nn*_-only fit (V1-V2 data; Fig. S9c). Comparing cross-validated EV_mod_ across nested fits attributes specific geometric components. Rather than testing all pairs, we test only comparisons answering one geometric question each. (i) Contributions of *α*_*sn*_ and *α*_*ns*_: the gap between the both-in-AIS and source-in-AIS (target-in-AIS) fits isolates *α*_*sn*_ (*α*_*ns*_). (ii) Their difference: source-in-AIS vs. target-in-AIS fits. (iii) Contribution of *α*_*nn*_: no pair of fits differs by *α*_*nn*_ alone, as the unconstrained fit adds *α*_*sn*_ + *α*_*nn*_ to the target-in-AIS fit and *α*_*ns*_ + *α*_*nn*_ to the source-in-AIS fit. If (i) detects no contribution from one of the two off-diagonal components, a non-significant gap for the comparison adding it together with *α*_*nn*_ indicates no detectable *α*_*nn*_ (ACA-VIP: unconstrained vs. target-in-AIS) - this reuses the unconstrained fit of the main analysis, requiring no additional model. If instead both off-diagonal components contribute (V1-V2), nested comparisons cannot isolate *α*_*nn*_, and we resort to the more computationally expensive option of fitting an *α*_*nn*_-only model and testing it against the residual-shuffle null (Appendix B.13). This yields four comparisons for ACA-VIP and three for V1-V2 data (Fig. 4d,h). Fig. S7e-h illustrates this for the four individual ground-truth geometries: when *α*_*ss*_ = 1 the both-in-AIS constraint captures the unconstrained EV_mod_; when *α*_*ns*_ = 1 (*α*_*sn*_ = 1) only the target-in-AIS (source-in-AIS) constrain captures the unconstrained EV_mod_, while constraining the source (target) reduces EV_mod_ to zero; when *α*_*nn*_ = 1 any constraint reduces EV_mod_ to zero. The *α* attribution from this approach is not univocal - nested gaps quantify the variance explained by adding a geometric component on top of a constrained baseline, rather than yielding a closed-form additive decomposition of EV_mod_ into the four *α* terms - but comparison between these constrained models suffices to localize which components carry modulatory variance.

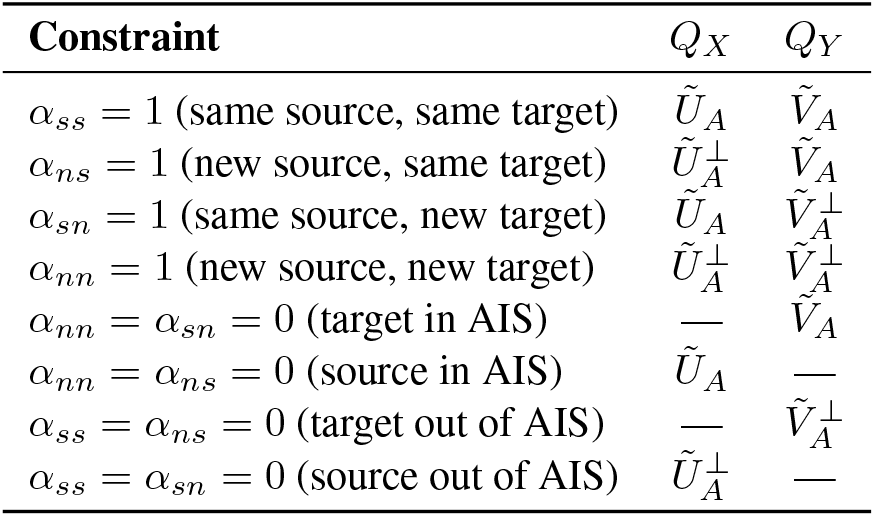

| **Constraint** | *Q*<sub>*X*</sub> | *Q*<sub>*Y*</sub> |
|---|---|---|
| *α*<sub>*ss*</sub> = 1 (same source, same target) | *Ũ*<sub>*A*</sub> | *Ṽ*<sub>*A*</sub> |
| *α*<sub>*ns*</sub> = 1 (new source, same target) | *Ũ*<sub>*A*</sub><sup>⊥</sup> | *Ṽ*<sub>*A*</sub> |
| *α*<sub>*sn*</sub> = 1 (same source, new target) | *Ũ*<sub>*A*</sub> | *Ṽ*<sub>*A*</sub><sup>⊥</sup> |
| *α*<sub>*nn*</sub> = 1 (new source, new target) | *Ũ*<sub>*A*</sub><sup>⊥</sup> | *Ṽ*<sub>*A*</sub><sup>⊥</sup> |
| *α*<sub>*nn*</sub> = *α*<sub>*sn*</sub> = 0 (target in AIS) | — | *Ṽ*<sub>*A*</sub> |
| *α*<sub>*nn*</sub> = *α*<sub>*ns*</sub> = 0 (source in AIS) | *Ũ*<sub>*A*</sub> | — |
| *α*<sub>*ss*</sub> = *α*<sub>*ns*</sub> = 0 (target out of AIS) | — | *Ṽ*<sub>*A*</sub><sup>⊥</sup> |
| *α*<sub>*ss*</sub> = *α*<sub>*sn*</sub> = 0 (source out of AIS) | *Ũ*<sub>*A*</sub><sup>⊥</sup> | — |

Here 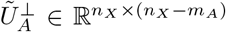 and 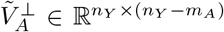 are orthonormal bases for the orthogonal complements of the baseline source and target AIS, and “—” indicates that the corresponding factor is left unconstrained. Constraining to the “same” subspaces projects the data onto *m*_*A*_-dimensional subspaces, making the ALS block systems much cheaper (block size drops from *n*_*X*_ × *n*_*X*_ to *m*_*A*_ × *m*_*A*_).

##### Remark B.6

(Constrained fitting vs. post-hoc *α* indices). The constrained model directly optimizes modulation within a specified geometric subspace, whereas the post-hoc *α* indices decompose an unconstrained solution. These can give different results: the unconstrained fit might place a channel at an intermediate angle (e.g., *α*_*ss*_ ≈ *α*_*ns*_ ≈ 0.5), whereas the constrained *α*_*ss*_ model finds the best purely within-subspace solution. Comparing the cross-validated EV_mod_ of constrained versus unconstrained models provides a direct test of whether modulatory variance is attributable to specific geometric components, complementing the descriptive *α* indices.

### B.9 Simulation details

In this section, we provide additional implementation details on all simulations of the paper. Simulations in Figs. 2, 3, S1, and S2, S3, S4, and S7 all share the generative structure

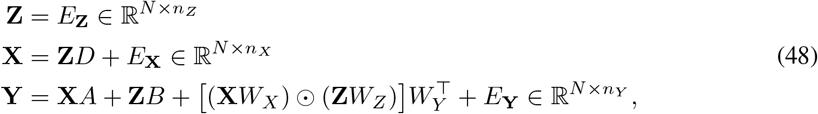

but differ in interaction structure (*A, B, D, W*_*X*_, *W*_*Y*_, *W*_*Z*_), population dimensionalities (*n*_*X*_, *n*_*Y*_, *n*_*Z*_), and sample size *N* . Noise terms *E*_**X**_, *E*_**Y**_, *E*_**Z**_ are i.i.d. Gaussian with unit variance unless stated otherwise. The low-rank factors are parametrized as

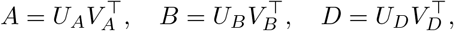

where the columns of 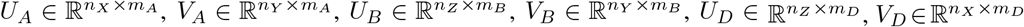, and the MIC weight vectors **w**_*X*_, **w**_*Y*_, **w**_*Z*_ are drawn independently and uniformly from the unit sphere, unless the simulation specifies a particular geometry. Throughout, rank selection uses the 1-SEM rule applied to 10-fold cross-validated EV curves.

#### Modulation strength sweep; Fig. 2(a-e)

We first considered a minimal scenario with *n*_*X*_ = 2, *n*_*Y*_ = 1, *n*_*Z*_ = 1, and *N* = 10^4^. The AIS matrix is fixed at *A* = [1 0]^*⊤*^, so that only the first source dimension drives *Y* additively (independent of *Z*), while the second drives *Y* multiplicatively through the MIC, with **w**_*X*_ = [0 1]^*⊤*^ · *γ*, **w**_*Y*_ = **w**_*Z*_ = 1. *m*_*B*_ = *m*_*D*_ = 0 throughout. Modulation strength *γ* is swept from 0 to 1.2 in steps of 0.2 over 5 independent repeats. AIS EV is additionally evaluated fitting the model on *X*_1_ and *X*_2_ separately, to make explicit which source component is captured by the AIS.

#### High-dimensional rank recovery; Fig. 2(f, g)

We then moved to a high-dimensional setting with *n*_*Z*_ = *n*_*X*_ = *n*_*Y*_ = 20 and *N* = 2000. Ground-truth ranks *m*_*A*_ and *m*_*C*_ are swept jointly from 0 to 5 (with *m*_*B*_ = *m*_*D*_ = 0 and *Z, X* drawn as independent Gaussians), with 5 independent simulations per rank pair, resulting in 30 simulations per marginal rank. Rank selection is followed by a permutation significance test on held-out predictions (1000 shuffles; *p <* 0.05 required to retain non-zero rank).

#### Data efficiency; Fig. 2h-i and Fig. S2

We assessed data efficiency by sweeping dimension *d* := *n*_*X*_ = *n*_*Y*_ = *n*_*Z*_ ∈ {10, 15, 20} and sample size *N* over 10 log-spaced values in [20, 20000], with *m*_*A*_ = 2, *m*_*C*_ = 1, *m*_*B*_ = *m*_*D*_ = 0, all factor matrices drawn uniformly on the unit sphere, and 30 simulations per (*d, N*). At each (*d, N*) the rank-selected fit (Algorithm 1) is compared against a full-rank counterpart in which the modulation tensor or AIS are fit without rank constraint. We report cross-validated AIS EV and EV_mod_ (Fig. 2h-i for *d* = 10; Fig. S2a-c, f-h, all *d*). To probe scaling, both EVs are then normalized by the ground-truth EV computed on each test fold from the true *A, W*_*X*_, *W*_*Y*_, *W*_*Z*_, and replotted against *N* rescaled by parameter count: *N/d* for low-rank fits, *N/d*^2^ for full-rank AIS, *N/d*^3^ for full-rank MICs (Fig. S2d-e, i-j).

#### Geometric decomposition, Fig. 3

We validated the geometric decomposition with *n*_*X*_ = 50, *n*_*Y*_ = 5, *n*_*Z*_ = 1, *m*_*A*_ = 2, *m*_*C*_ = 1, *N* = 1000, and 30 simulations per scenario. The source MIC weight **w**_*X*_ is drawn at a fixed angle *θ*_*X*_ to span(*U*_*A*_), while the target weight **w**_*Y*_ is always drawn within span(*V*_*A*_) (*θ*_*Y*_ = 0). Two ground-truth geometries are tested: *θ*_*X*_ = 0, corresponding to 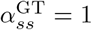 with all other components zero (Fig. 3b-d), and 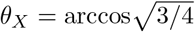, corresponding to 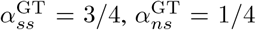 (Fig. 3e-g). The shuffle null for the *α* indices is constructed from 1000 random draws of (**w**_*X*_, **w**_*Y*_) conditioned on the fitted *Â*, and constrained-fit EV_mod_ is evaluated for all combinations of source and target geometric constraints.

**Figure S1:**
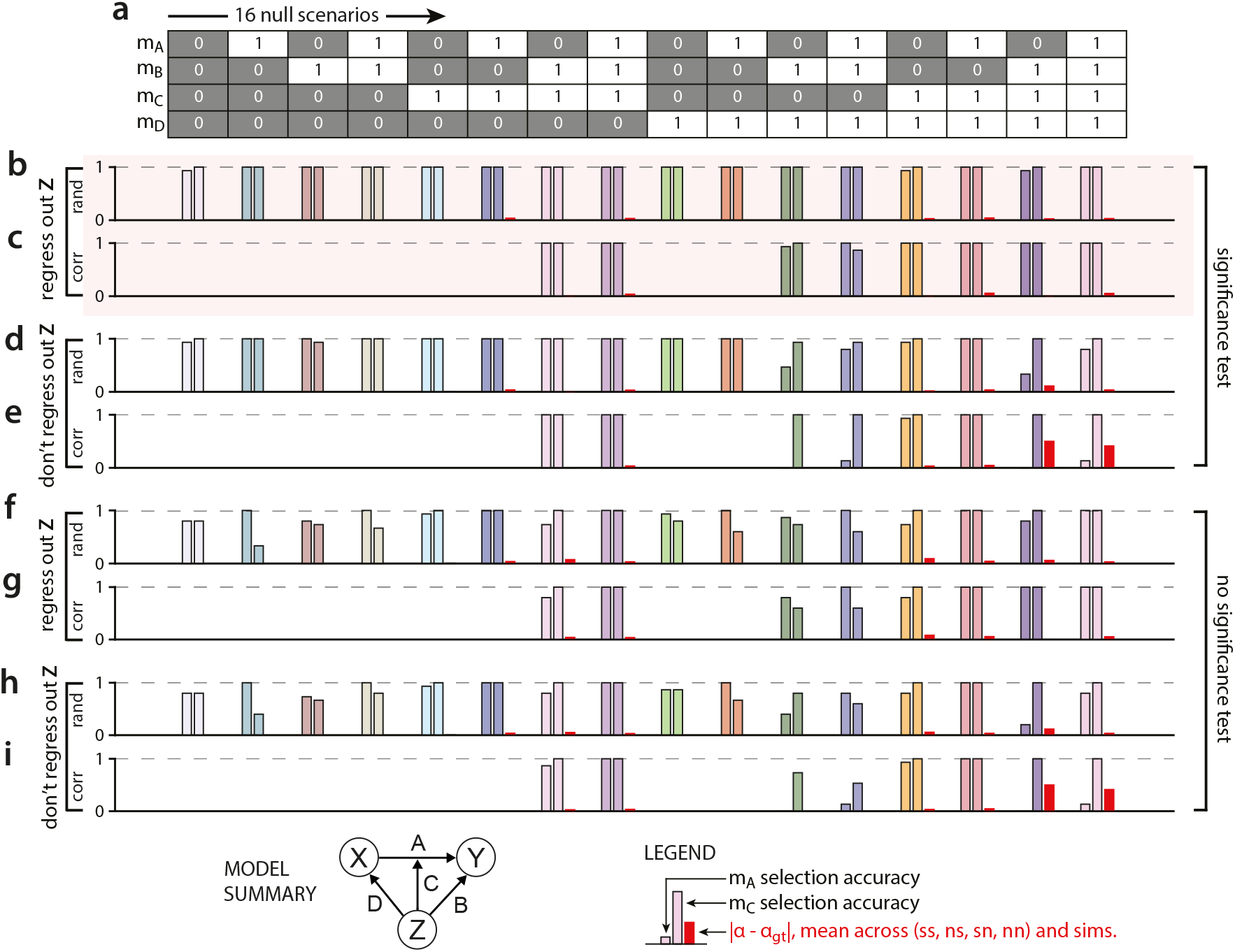
Pipeline validation across all 16 rank combinations. (a) Ground-truth rank indicator matrix: *m*_*A*_, *m*_*B*_, *m*_*C*_, *m*_*D*_∈ {0, 1} for each of the 16 simulated scenarios (columns). The model schematic (bottom left) is reproduced from Fig. 1b for reference. Panels (b-i) evaluate the MICs fitting pipeline across a 2 × 2 × 2 factorial design: whether *Z* is regressed out of *X* and *Y* (Step 1 of Algorithm 1; panels b–c, f–g vs. d–e, h–i), whether the shuffle significance test defined in App. B.13 is applied to EV_mod_ (panels b–e vs. f–i), and whether ground-truth interaction subspace vectors are drawn uniformly from the unit sphere (*rand*) or set to be perfectly co-aligned across model terms (*corr*; see below). In each sub-panel, bars for each scenario show *m*_*A*_ selection accuracy (left bar), *m*_*C*_ selection accuracy (center bar), and mean absolute geometry index error, averaged over the four geometric components (ss, ns, sn, nn) and simulations (right bar, red). (b) Pipeline with Step 1 and permutation significance test (*rand* condition): ground-truth weight vectors are drawn independently and uniformly from the unit sphere. (c) Same as (b), *corr* condition: for scenarios in which at least two of *m*_*B*_, *m*_*C*_, *m*_*D*_ are non-zero, the relevant low-rank components are set to be perfectly co-aligned. Specifically, when *m*_*B*_ = *m*_*C*_ = 1, the MIC and the *Z* − *Y* interaction subspace are perfectly aligned in both *Z* and *Y* ; when *m*_*D*_ = *m*_*C*_ = 1, the MIC and the *Z* − *X* interaction subspace are perfectly aligned in both *Z* and *X*; when *m*_*B*_ = *m*_*D*_ = 1, the *Z* → *Y* and *Z* → *X* interaction subspaces are perfectly aligned in *Z*. (d-e) Same as (b-c) but without regressing *Z* out of *X* and *Y* (Step 1 omitted). (f-g) Same as (b-c) but without the permutation significance test. (h-i) Same as (b-c) but without Step 1 and without the permutation significance test. We ran 15 independent simulations per scenario, with *N* = 1000 samples per simulation and *n*_*X*_ = *n*_*Y*_ = *n*_*Z*_ = 5 in all simulations.

**Figure S2:**
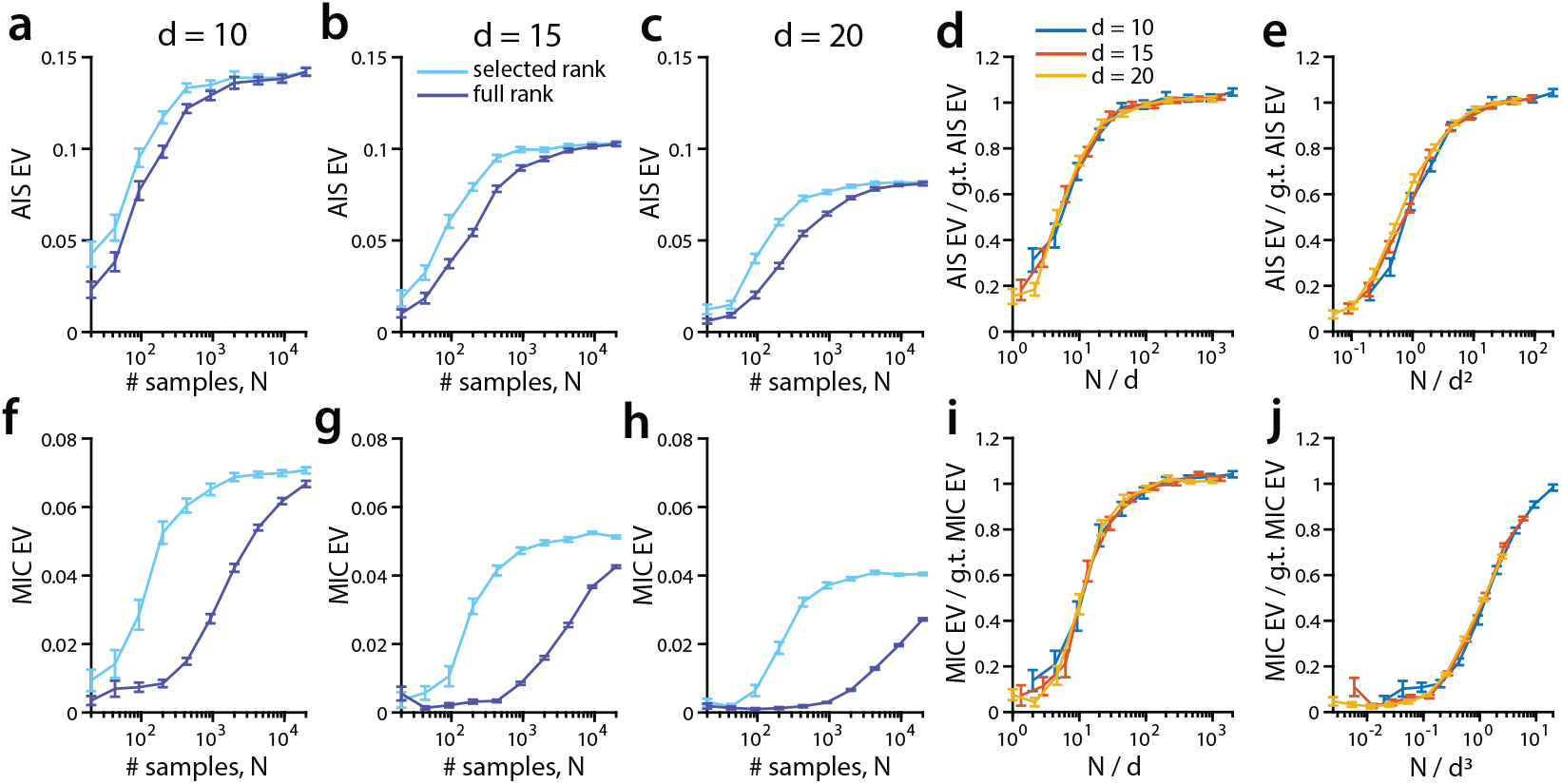
Data efficiency of MICs relative to full-rank models. (a) Variance explained by AIS as a function of sample size *N*, for the rank-selected low-rank model (cyan) and the full-rank model (blue), with *n*_*X*_ = *n*_*Y*_ = *n*_*Z*_ = 10 =: *d*. Lines show mean ± SEM across 30 independent simulations. (b-c) Same as (a), with *d* = 15 and *d* = 20. (d) Scaling collapse of low-rank AIS explained variance for the three values of *d* in (a-c). The number of samples is rescaled by a factor proportional to the number of parameters in the low-rank AIS model (specifically, 1*/d*) and the explained variance is rescaled by the explained variance of the ground-truth AIS. (e) Same as (d), for the full-rank AIS model (∝ *d*^2^ parameters). (f-h) Same as (a-c), for the low-rank and full rank MIC models. (i-j) Same as (d-e) for the low-rank (number of parameters ∝ *d*) and full-rank (number of parameters ∝ *d*^3^) MIC models. Here the explained variance is rescaled by the explained variance of the ground-truth MIC. In all simulations, *m*_*A*_ = 2 and *m*_*C*_ = 1.

#### Models sweep with pipeline ablations; Fig. S1

We performed an exhaustive sweep over all 2^4^ = 16 binary rank combinations (*m*_*A*_, *m*_*B*_, *m*_*C*_, *m*_*D*_ ∈ {0, 1}) with *n*_*Z*_ = *n*_*X*_ = *n*_*Y*_ = 5, *N* = 1000, and 15 simulations per scenario. For each scenario, two subspace alignment conditions are tested: *rand*, in which all weight vectors are drawn independently from the unit sphere; and *corr*, applied only when at least two of *m*_*B*_, *m*_*C*_, *m*_*D*_ are non-zero, in which the relevant weight vectors of all non-zero terms are set equal to a shared unit vector per space, making all interaction subspaces perfectly co-aligned. The full 2 × 2 × 2 factorial of pipeline conditions - (regress out *Z*) × (permutation significance test) × (rand/corr) - is evaluated, yielding panels (b-i) of Fig. S1. The significance test uses 1000 permutation shuffles.

#### Non-Gaussian emission models, Fig. S3

This simulation departs from the others only in the observation step. To test robustness to emission-model misspecification, we used the generative process of Eq. 48 as a latent rate-level model and observed counts through a non-linear emission. Specifically, the latent variables **X, Y** produced by Eq. 48 model, with *W*_*Y*_ rescaled by a modulation-strength scalar *γ*, are interpreted as log-rates after a constant baseline shift: 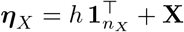 and 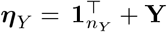, with *h* = 2 and entrywise clipping |*η* | ≤ 4 for numerical stability. Three emission models were compared:

- *Gaussian benchmark*: **X**^obs^ = **X, Y**^obs^ = **Y** (the latent rate-level variables, recovering the standard setup);
- *Poisson*: 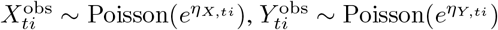, independently across entries;
- *zero-inflated Poisson* (zero-inflation probability *p* = 0.5): 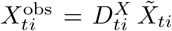 with 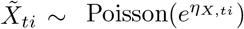 and 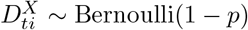, analogously for *Y* .

**Figure S3:**
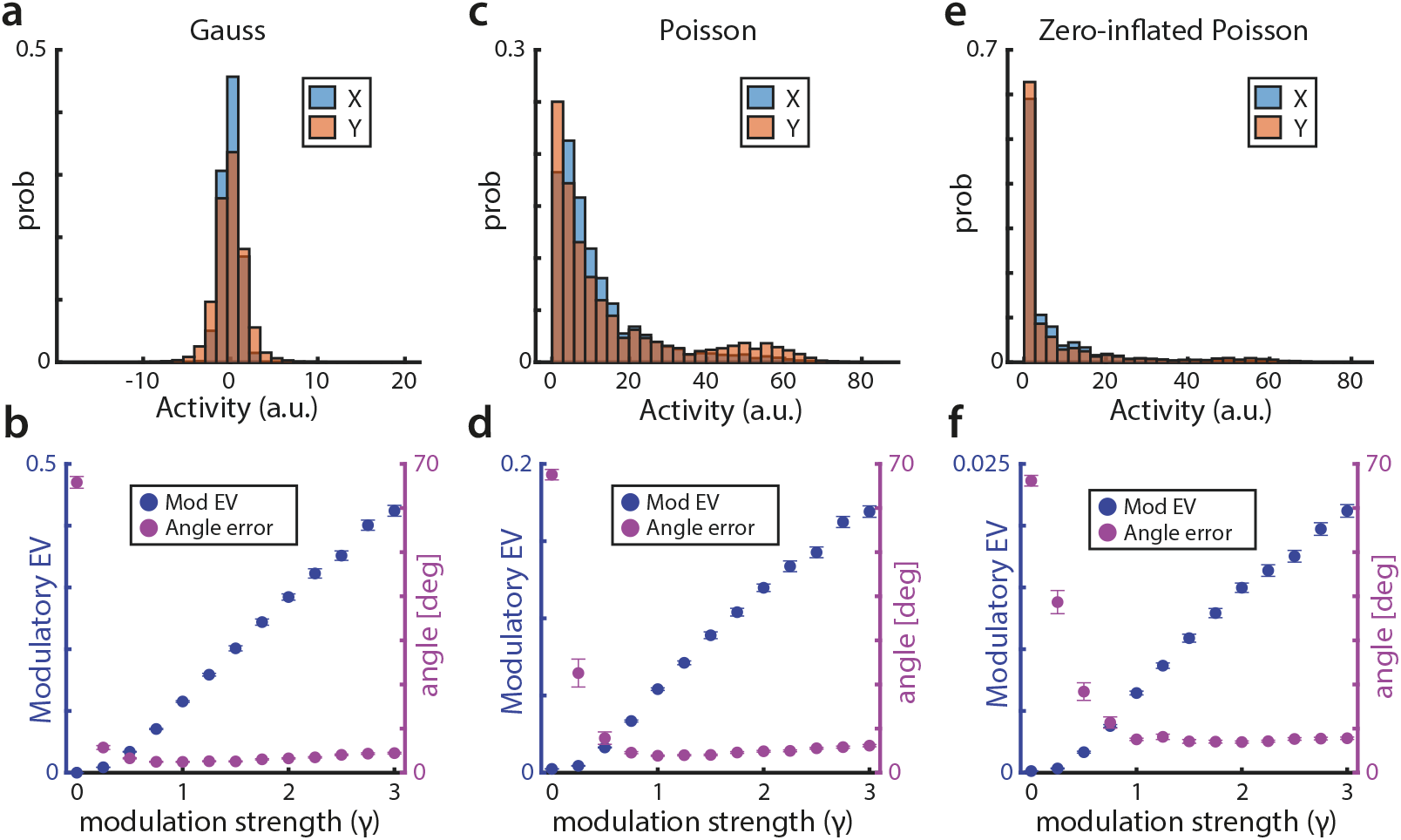
MICs performance under non-Gaussian observation models. All conditions share the same latent bilinear generative model, only the emission distribution of *X* and *Y* varies across scenarios. (a) Marginal distribution of *X* and *Y* for one example simulation in the Gaussian (linear benchmark) condition. (b) EV_mod_ (blue) and mean angle between estimated and ground-truth MIC weight vectors (purple) as a function of modulation strength *γ*. (c-d) Same as (a-b) for Poisson-distributed *X* and *Y* . (e-f) Same as (a-b) for zero-inflated Poisson observations (zero-inflation probability *p* = 0.5). *n*_*X*_ = *n*_*Y*_ = *n*_*Z*_ = 5; ground-truth ranks *m*_*A*_ = *m*_*B*_ = *m*_*C*_ = 1, *m*_*D*_ = 0; *N* = 10,000 samples per simulation. Curves show mean ± SEM across 50 independent simulations.

We swept *γ* ∈ [0, 3] in steps of 0.25, with 50 simulations per *γ*. Across conditions, *n*_*X*_ = *n*_*Y*_ = *n*_*Z*_ = 5, *N* = 10^4^, *m*_*A*_ = *m*_*B*_ = *m*_*C*_ = 1, *m*_*D*_ = 0. The MICs pipeline – which assumes Gaussian observations – was applied without modification, fitting at the ground-truth ranks. We report EV_mod_ and the mean principal angle between fitted and ground-truth (**w**_*X*_, **w**_*Y*_, **w**_*Z*_) as a function of *γ*. EV_mod_ increased monotonically with *γ* in all three conditions Fig. S3b,d,f), and the mean angle error decreased toward zero as *γ* grew, indicating reliable recovery of the ground-truth MIC geometry despite emission-model misspecification. As expected, absolute EV_mod_ was attenuated under count noise relative to the Gaussian benchmark, with zero-inflated Poisson showing the largest attenuation.

#### Comparison with *Z*-splitting baseline; Fig. S4a-b

To benchmark MICs against the Z-splitting baseline, we simulated data with *n*_*X*_ = *n*_*Y*_ = 5, *n*_*Z*_ ∈ {1, 3, 5, 7, 9}, *N* = 500, *m*_*A*_ = 2, *m*_*C*_ = 1, and *m*_*B*_ = *m*_*D*_ = 0, running 100 independent simulations per *n*_*X*_ value. For the splitting baseline, **Z** is projected onto its first principal component and samples are partitioned into *N*_bins_ ∈ {1, …, 15} equipopulated quantile bins; a separate rank-selected AIS is fit within each bin using the same cross-validation partition as MICs, and modulatory EV is defined as the difference between the mean within-bin AIS EV and the EV of a single AIS fit pooled across all samples. For *n*_*Z*_ *>* 1, this projection discards variance in **Z** beyond its leading principal component. MIC modulatory EV and the peak splitting EV (maximized over *N*_bins_) are then compared as a function of *n*_*Z*_.

#### Single-component and multi-MIC geometries, Fig. S7

We extended the geometric-decomposition validation in two directions. Across all panels, *n*_*X*_ = *n*_*Y*_ = 20, *n*_*Z*_ = 10, *m*_*A*_ = 2, *m*_*B*_ = *m*_*D*_ = 0, and *N* = 1000. Panels (a-h) cover the four pure single-component scenarios 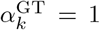, *k*∈ {*ss, ns, sn, nn*}, with *m*_*C*_ = 1 and 10 simulations per scenario: **w**_*X*_ is drawn uniformly on the unit sphere of either span(*U*_*A*_) (*s* on the source side) or its orthogonal complement (*n*), and analogously for **w**_*Y*_ with respect to span(*V*_*A*_); **w**_*Z*_ is drawn uniformly on the unit sphere of 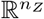. For each fitted MIC, null-subtracted *α* indices use 1000 random-rotation shuffles, and constrained-fit EV_mod_ is evaluated under all four combinations of source and target geometric constraints (no/source/target/both). Panels (i-k) cover multi-MIC scenarios with *m*_*C*_ ∈ { 1, 2, 3} and 30 simulations per *m*_*C*_: for each channel, angles *θ*_*X*_, *θ*_*Y*_ are drawn independently and uniformly from [0, *π/*2], and **w**_*X*_, **w**_*Y*_ are constructed at these angles to span(*U*_*A*_) and span(*V*_*A*_) respectively, yielding randomly distributed ground-truth indices 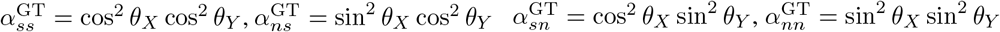.

**Figure S4:**
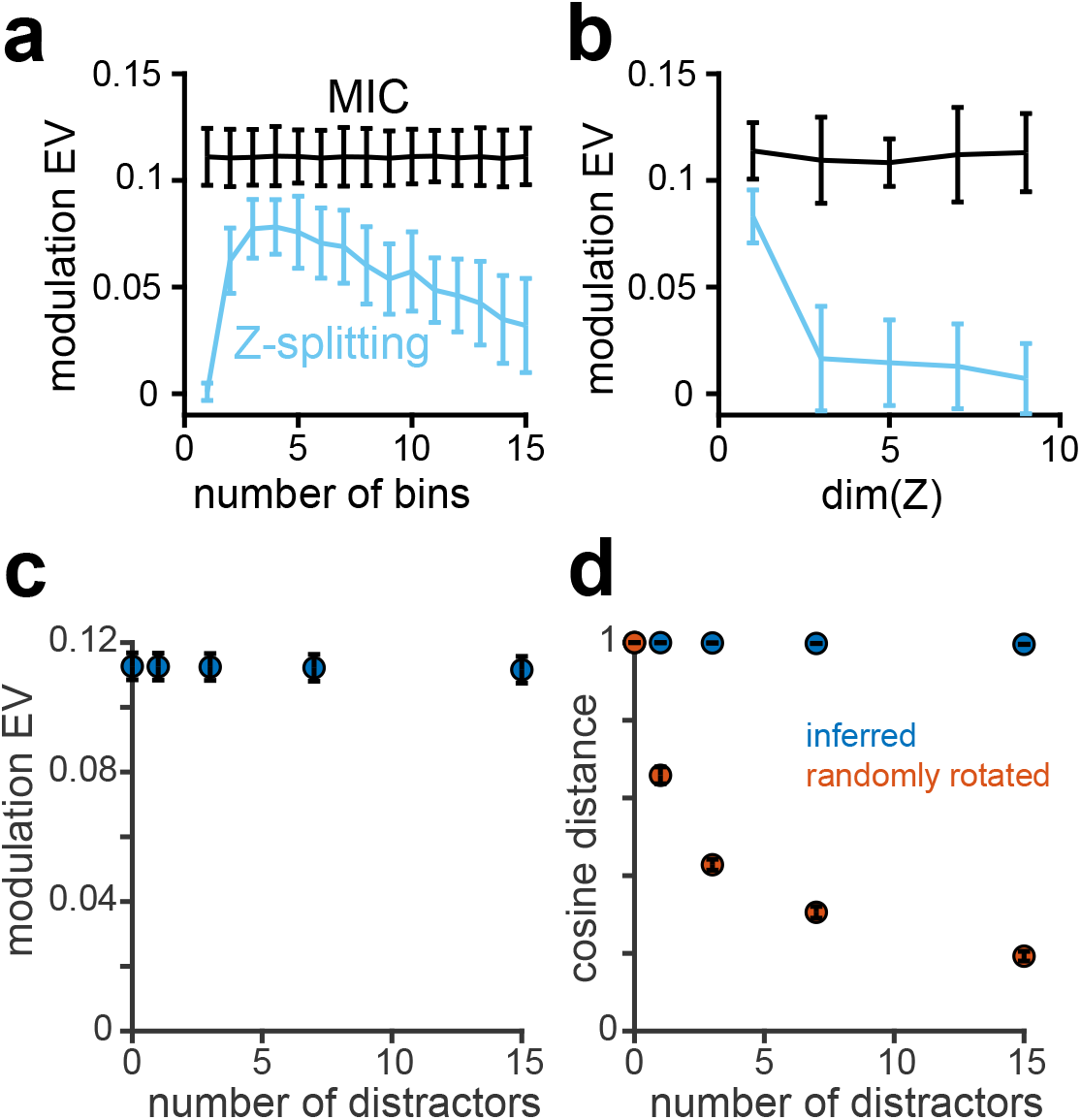
Benchmarking MICs versus Z-splitting. (a) Modulatory EV recovered by the MICs method (black) versus a benchmark that discretizes a 1D *Z* into quantile bins and refits the AIS within each bin (light blue), as a function of the number of bins. (b) Same comparison as a function of dim(*Z*), with the quantile-splitting benchmark applied to the first principal component of *Z*; 100 simulations per scenario, *N* = 500 samples per simulation. (c) EV_mod_ recovered by MICs when one modulating dimension of *Z* is embedded among *n* non-modulatory distractor dimensions (*n*_*Z*_ = *n* + 1), as a function of *n*. (d) Cosine distance between ground-truth *w*_*Z*_ and the inferred *w*_*Z*_ (inferred) or a uniformly random unit vector in 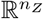 (randomly rotated). 20 simulations per *n, N* = 2000, *n*_*X*_ = *n*_*Y*_ = 5, *m*_*A*_ = 2, *m*_*C*_ = 1, *m*_*B*_ = *m*_*D*_ = 0.

#### Identifying components of *Z* that drive modulation; Fig. S4c

We embedded one genuinely modulatory variable within a pool of *n* non-modulatory “distractors” and asked whether MICs recovers it. We ran 20 independent simulations per value of *n* ∈ {1, 3, 7, 15 }, with *n*_*X*_ = *n*_*Y*_ = 5, *n*_*Z*_ = *n* + 1 (the *n* distractors plus one genuine), *N* = 2000, *m*_*A*_ = 2, *m*_*B*_ = *m*_*D*_ = 0, and *W*_*Z*_ = (1, 0, …, 0)^*⊤*^. As the number of distractors increased, EV_mod_ was essentially unchanged and the ground-truth and inferred *W*_*Z*_ remained closely aligned, indicating that MICs correctly recovered the genuine modulator.

In the ACA-VIP dataset, refitting MICs with wheel speed as the only modulator (i.e., removing pupil diameter from *Z*) did not reduce EV_mod_ (0.0066 ± 0.0011 wheel only vs. 0.0071 ± 0.0012 with both variables, *p* = 0.64; Fig. S11f), whereas refitting with pupil diameter as the only modulator significantly reduced it (0.0013 ± 0.0006; *p* = 0.0007 vs. both, *p* = 0.003 vs. wheel only). Thus, wheel speed is the main driver of the modulatory effect we observed.

**Figure S5:**
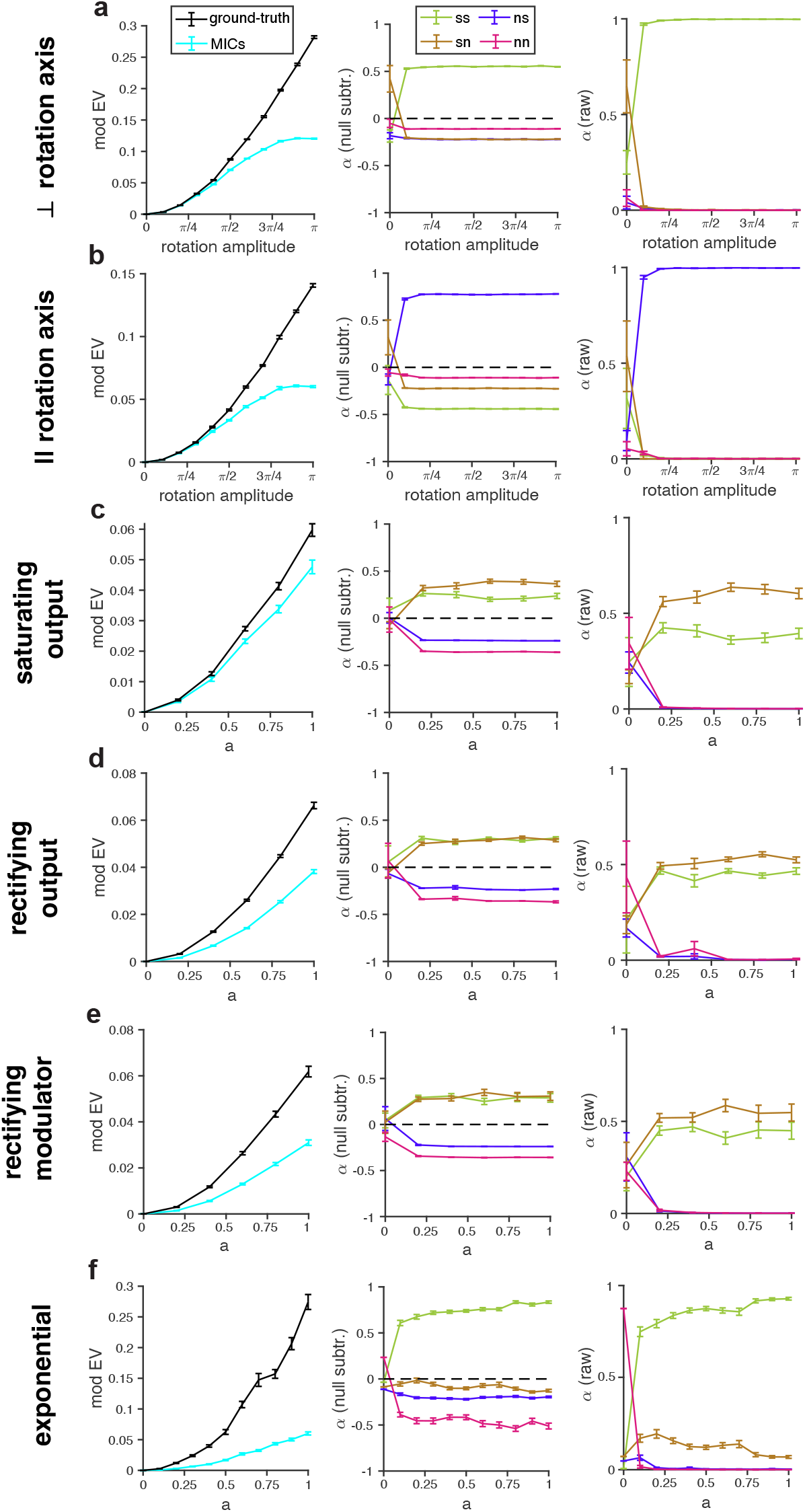
MICs applied to nonlinear generators. (a) Perpendicular-axis rotation generator (Appendidx B.10).Top: EV_mod_ recovered by MICs (cyan) and by the ground-truth generator (black) as a function of the rotation amplitude. Middle: null-subtracted geometry indices of the inferred MICs (*α*_*ss*_: green, *α*_*ns*_: blue, *α*_*sn*_: brown, *α*_*nn*_: magenta). Bottom: raw geometry indices. (b) Same as (a) for the parallel-axis rotation generator. (c-f) Same as (a) for the saturating output (c), rectifying output (d), rectifying modulator (e), and exponential (f) generators, as a function of the modulation strength parameter *a*. Mean ± SEM across 20 simulations per condition (10 for rotations), *N* = 10^4^ trials per simulation.

**Figure S6:**
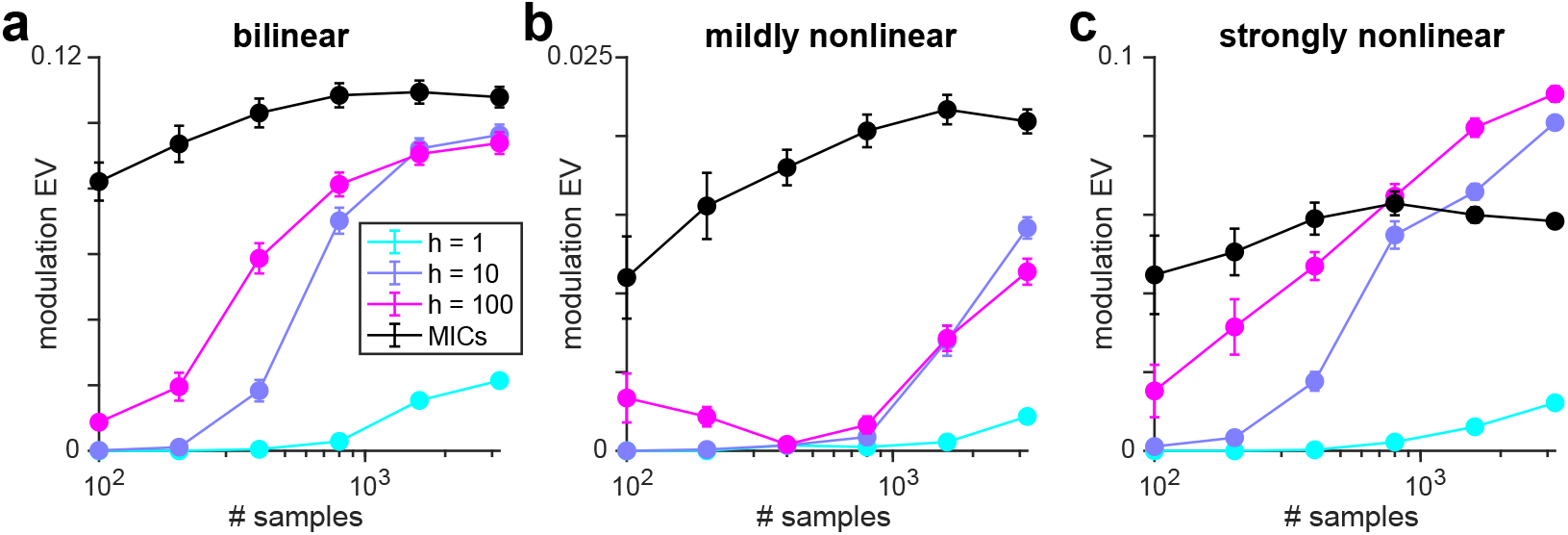
Sample efficiency of MICs versus a multilayer perceptron. (a) Cross-validated modulatory EV as a function of sample size *N* under a bilinear generator, for MICs and for one-hidden-layer MLPs with *h* · *m*_*C*_ hidden ReLU units. All models were fit to the residuals of the joint additive model using the same cross-validation folds (Appendix B.12). (b) Same as (a) for a mildly nonlinear generator (parallel-axis rotation, *a* = 2*π/*5; Appendix B.10). (c) Same as (a) for a strongly nonlinear generator (parallel-axis rotation, *a* = *π*). Mean ± SEM across 10 simulations per *N* .

### B.10 Stress-testing MICs on nonlinear generators

The MICs model (Eq. 5) assumes that the source-target interaction is linear and that modulation is bilinear. We applied MICs to simulations of five different classes of nonlinear generators that break these assumptions, all of the form

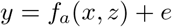

where *e* is zero-centered, unit-variance Gaussian noise and *a* is a scalar that controls the amount of modulation.

- *Rotations: f*_*a*_(*x, z*) = *A*^*⊤*^*R*(*θ* = *az*, **u**)*x*. Here, *n*_*X*_ = *n*_*Y*_ = 3, *n*_*Z*_ = 1, and *R*(*θ*, **u**) is the matrix that rotates vectors by angle *θ* about axis **u** in ℝ ^3^. For any fixed value of *z*, the source-target interaction is linear and characterized by an AIS (as assumed by MICs). Changes in *z*, however, elicit nonlinear changes in the AIS, breaking the assumptions of MICs. In particular, the AIS rotates on the source side. We set 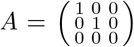 and either **u** = (0, 0, 1)^*⊤*^ (“perpendicular”; the source-side AIS rotates within itself) or **u** uniformly drawn from span (1, 0, 0)^*⊤*^, (0, 1, 0)^*⊤*^ (“parallel”; the source-side AIS rotates outside itself). Increasing *a* increases the rotation amplitude and hence the amount of modulation by *z*. We ran 10 independent simulations per value of *a* and type of rotation, with *N* = 10^4^ trials per simulation and the components of *x* and *z* drawn independently from the zero-centered Gaussian with unit variance on each trial.
- *Saturating output* 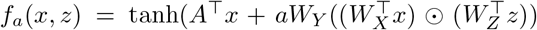. If tanh were removed, the remaining expression would simply be the MICs model. We set *n*_*X*_ = *n*_*Y*_ = 5, *n*_*Z*_ = 1, *m*_*A*_ = 2, *m*_*C*_ = 1, 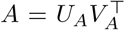 with the columns of 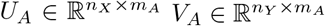 drawn uniformly from the unit sphere, 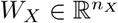 drawn uniformly from the column space of *U*_*A*_, 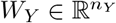 drawn uniformly from the subspace making an angle *π/*4 to the column space of *V*_*A*_, and *W*_*Z*_ = 1. This setup ensures that the MICs model embedded within the output nonlinearity has geometry indices *α*_*ss*_ = *α*_*sn*_ = 1*/*2. As before, *a* controls the amount of modulation by *z*. We ran 20 simulations per value of *a*, again with *N* = 10^4^ trials and *x*_*i*_, *z*_*j*_ ∼ *N* (0, 1) on each trial.
- *Rectifying output:* 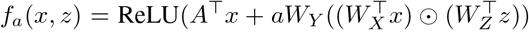. All other details are the same as in the *Saturating output* case.
- *Rectifying modulator:* 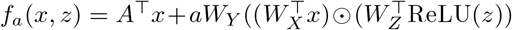. All other details are the same as in the *Saturating output* case.
- *Exponential: f*_*a*_(*x, z*) = (*A*^*⊤*^*x*)^1+*a*(2*z−*1)^. The exponentiation acts componentwise. We set *n*_*X*_ = *n*_*Y*_ = 5, *n*_*Z*_ = 1, *m*_*A*_ = 2, and 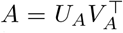 with the columns of 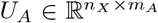 drawn uniformly from the portion of the unit sphere where all components are positive. If *a >* 0, then *z >* 1*/*2 results in a supralinear mapping from *x* to *y, z <* 1*/*2 in a sublinear mapping, and *z* = 1*/*2 in a linear mapping. In each trial (20 simulations per *a*, each with *N* = 10^4^ trials), we drew the components of *x* and *z* uniformly from the interval (0, 1). Thus, the generator is well-defined for |*a*| ≤ 1 (for |*a*| *>* 1, the exponent can become negative).

**Figure S7:**
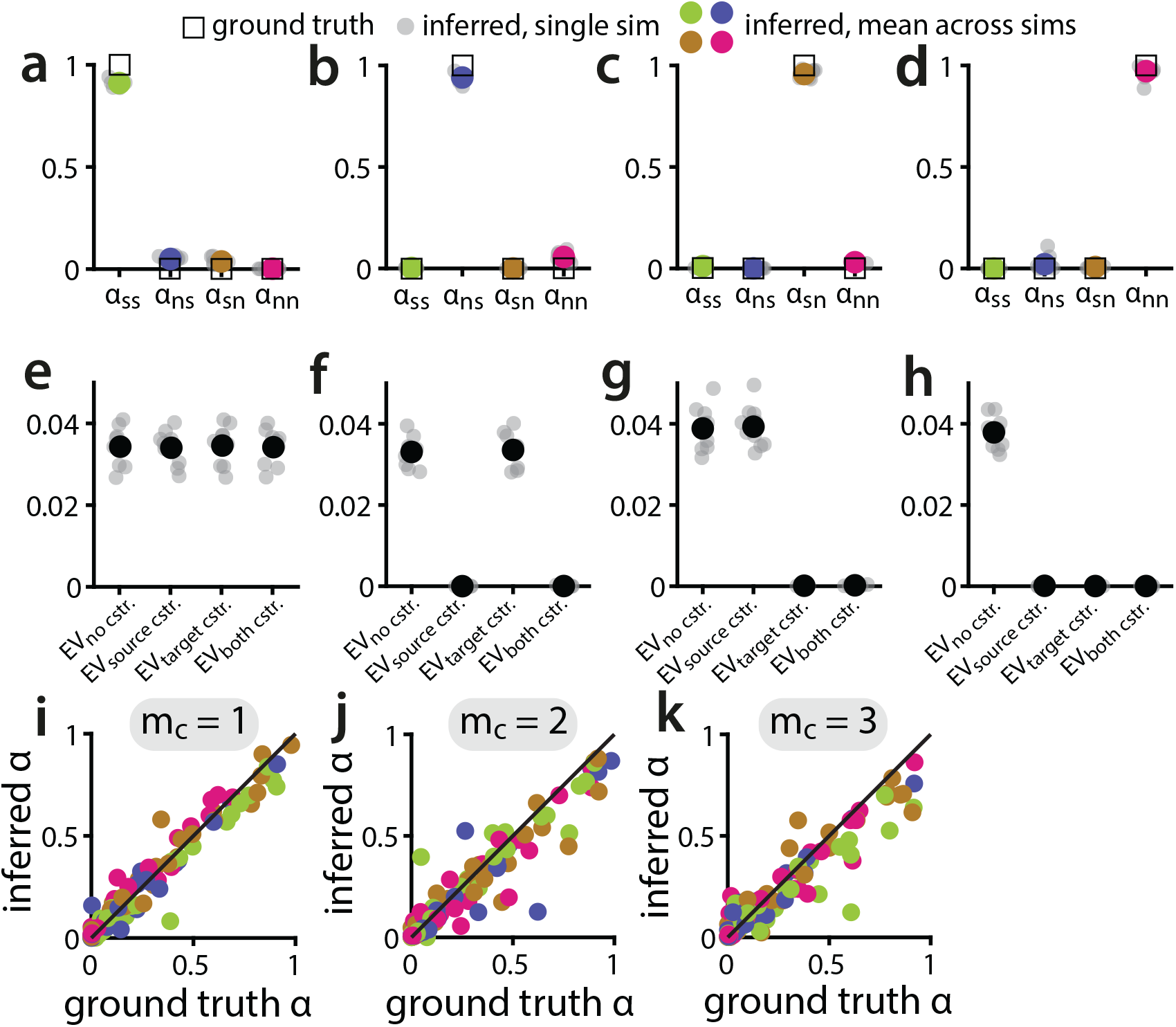
Validating the geometric decomposition across single-component and multi-MIC scenarios. (a) Raw geometry indices for a single-MIC scenario with ground-truth 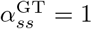, all other components zero. Open squares: ground truth; filled colored circles: mean across simulations; light grey dots: individual simulations. (b-d) Same as (a) for ground-truth 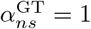(b), 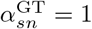 (c), and 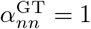 (d). (e) Constrained-fit EV_mod_ for the 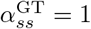 scenario, evaluated under four constraints: *no cstr*., unconstrained fit; *source cstr*., source factor confined AIS, forcing *α*_*ns*_ = *α*_*nn*_ = 0; *target cstr*., target confined to AIS, forcing *α*_*sn*_ = *α*_*nn*_ = 0; *both cstr*., both source and target confined to AIS, leaving only *α*_*ss*_ admissible. Filled black circles: mean across simulations; light grey dots: individual simulations. (f-h) Same as (e) for the 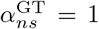 (f), 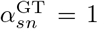 (g), and 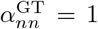 (h) scenarios. (i-k) Inferred against ground-truth *α* indices in multi-MIC scenarios with *m*_*C*_ = 1 (i), *m*_*C*_ = 2 (j), *m*_*C*_ = 3 (k); for each MIC, **w**_*X*_ is drawn to yield randomly-distributed ground-truth *α* profiles. Each dot is one channel from one simulation; colors indicate the four geometric components (*α*_*ss*_: green, *α*_*ns*_: blue, *α*_*sn*_: brown, *α*_*nn*_: magenta). Black line: identity. We ran 10 simulations per scenario in (a-h) and 30 simulations per *m*_*C*_ in (i-k). *n*_*X*_ = *n*_*Y*_ = 20, *n*_*Z*_ = 10, *m*_*A*_ = 2, *m*_*B*_ = *m*_*D*_ = 0, *N* = 1000 in all simulations.

**Figure S8:**
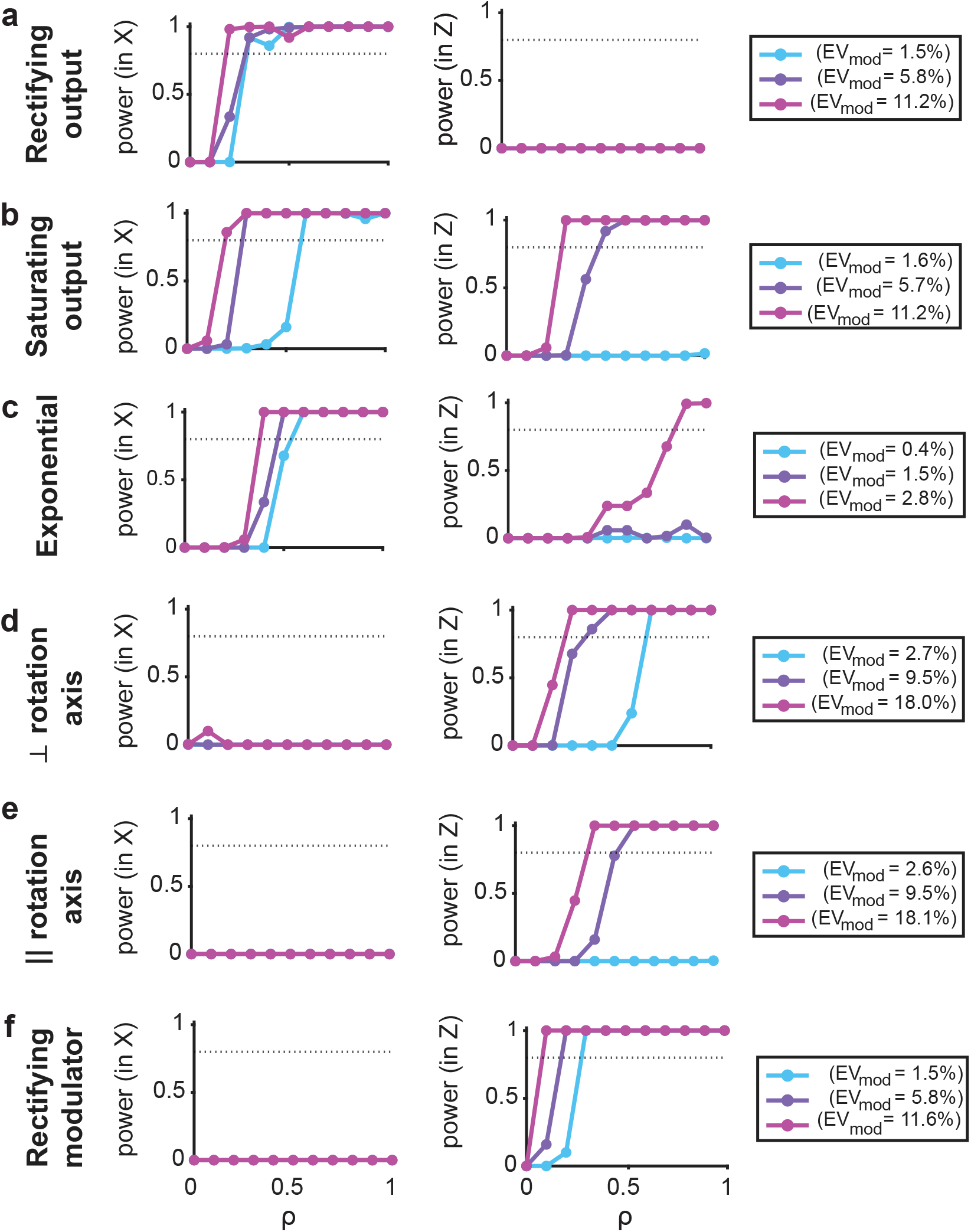
Power of the bilinearity diagnostic. Each nonlinear generator of Appendix B.10 was linearly interpolated with a bilinear MICs generator, so that *ρ* = 0 is purely bilinear and *ρ* = 1 is the original nonlinear generator. (a) Rectifying output generator. Left: power of detecting ΔEV_NL_ *>* 0 with the nonlinear-in-*x*. Right: power of detecting ΔEV_NL_ *>* 0 with nonlinear-in-*z* (sign test across 50 simulations, *α* = 0.05; dotted line: power of 0.8), as a function of *ρ*. Colors: modulation strength *a* = 1*/*3 (light blue), 2*/*3 (violet) and 1 (purple); legends report the EV_mod_ recovered by MICs at *ρ* = 0 for each *a*. (b) Same as (a) for the saturating output generator. (c) Same as (a) for the exponential generator. (d-e) Same as (a) for the perpendicular-axis (d) and parallel-axis (e) rotation generators (rotation amplitude *aπ*). (f) Same as (a) for the rectifying modulator generator. *N* = 2 × 10^4^ trials per simulation.

We used two criteria to assess the performance of MICs on these nonlinear generators. First, we compared the EV_mod_ returned by MICs to the “ground-truth” EV_mod_, defined as the explained variance of the ground-truth generator’s deterministic part, minus that of the same generator with *z* clamped to baseline (*z* = 1*/*2 for the *Exponential* generator, and *z* = 0 for all others). Across all generators, the EV_mod_ of MICs monotonically increased with the ground-truth value as *a* increased, indicating that MICs could still track changes in the amount of modulation despite the generator mismatch (Fig. S5).

Second, we asked whether the modulation geometry indices inferred by MICs match the geometry built into the ground-truth generators. The *Rotations* generator alters which source (*x*) patterns interact with a fixed set of target (*y*) patterns, and hence should have *α*_*sn*_ ≈ *α*_*nn*_ ≈ 0. In the perpendicular axis case, the rotation acts within the AIS, so we expect *α*_*ss*_ ≈ 1. In the parallel axis case, on the other hand, the rotation acts outside the AIS, so we expect *α*_*ns*_ ≈ 1. MICs recovered all of these expectations, regardless of the rotation amplitude (Fig. (Fig. S5)a-b).

The *Saturating output, Rectifying ouput*, and *Rectifying modulator* generators were constructed so that, if the nonlinearity were removed, the ground-truth geometry indices would be *α*_*ss*_ = *α*_*sn*_ = 1*/*2 and *α*_*ns*_ = *α*_*nn*_ = 0. Despite the nonlinearities, MICs identified *α*_*ss*_ and *α*_*sn*_ as the indices exceeding the random rotation null (Fig. S5c-e). For the two output nonlinearities, *x* enters only through 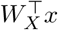 and *A*^*⊤*^*x*, both within the source AIS, so *α*_*ns*_ = *α*_*nn*_ = 0 is preserved, whereas the nonlinearity acts on the target and distorts the target MIC axis out of the target AIS, tilting the indices toward *α*_*sn*_ *> α*_*ss*_.

Under the *Exponential* generator, *z* controls the exponent applied to the baseline output *A*^*⊤*^*x*. Thus, *z* primarily changes the overall gain of *x*-to-*y* interactions but can also distort which patterns in *y* are targeted by *x*. Accordingly, MICs returns *α*_*ss*_ as the largest index, with a smaller (but above random-rotation null) *α*_*sn*_ (Fig. (Fig. S5)f).

### B.11 Diagnosing violations of bilinear modulation assumption

The MICs model assumes that modulation is bilinear, i.e. linear in the modulator *z* at fixed *x* and linear in the source *x* at fixed *z*. This assumption is deliberate: bilinearity, together with the low-rank CP parameterization, is what makes MICs data-efficient and geometrically interpretable. To diagnose violations of bilinearity, we added higher-order nonlinear terms to MICs and measured the change in performance compared to the bilinear baseline. Specifically, for a fitted channel (*w*_*X*_, *w*_*Y*_, *w*_*Z*_), define the residualized source projection *s*_*X*_ = **X**^*′*^*w*_*X*_, the modulator projection *s*_*Z*_ = **Z***w*_*Z*_, and the modulation-residual target projection *s*_*Y*_ = **Y**^*′′*^*w*_*Y*_ . We tested the performance of a nested set of regressions of *s*_*Y*_ :

- *Bilinear: s*_*Y*_ ∼ *s*_*X*_ ⊙ *s*_*Z*_.
- *Nonlinear in x:* 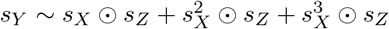.
- *Nonlinear in z:*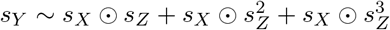.

We included terms up to third order to capture both rectification (second order) and saturation (third order) effects. We use the same cross-validation folds as were used to fit the MICs model and calculate the projections (*s*_*X*_, *s*_*Y*_, *s*_*Z*_) for the train and test sets using the axes (*w*_*X*_, *w*_*Y*_, *w*_*Z*_) from the train set. Each regression is ridge-regularized, with the penalty optimized by cross-validation over [10^*−*4^, 10^1^]. We report the difference ΔEV_NL_(*x*) (ΔEV_NL_(*z*)) between the cross-validated explained variance of the *Nonlinear in x* (*Nonlinear in z*) regression and the bilinear regression. ΔEV_NL_(*x*) *>* 0 indicates a violation of linearity in *x*, and similarly ΔEV_NL_(*z*) *>* 0 indicates a violation in *z*. Significance across simulation runs (or sessions, for real data analyses S9e and S11g) was assessed with a one-tailed sign (binomial) test on ΔEV_NL_ *>* 0, one-tailed because only an increase in EV indicates a violation of bilinearity. In simulations, we report power: the probability that the sign test detects nonlinearity (*p <* 0.05) across *n* = 50 runs, computed from the fraction of runs with ΔEV_NL_ *>* 0. For the bilinear generator (*ρ* = 0), this is the false-positive rate (Fig. S8).

We tested the diagnostic on simulations of the nonlinear generators in Appendix B.10 (Fig. S8). In the *Rotations* and *Rectifying modulator* generators, the nonlinearity acts directly on *z*, so the diagnostic should report a violation in *z* only. To determine how the diagnostic should behave on the remaining generators, we expanded them in power series around the baseline values of *x* and *z*:

- *Saturating output:* Besides the bilinear term, the lowest-order nonzero modulatory (i.e. containing both *x* and *z*) term is 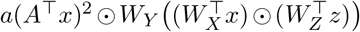, where the exponentiation acts componentwise. Thus, we expect the diagnostic to localize the violation primarily to *x*.
- *Rectifying output:* ReLU is not differentiable. However, it can be approximated by the smooth softplus function, ReLU 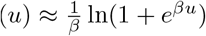, with the error vanishing as *β → ∞*. Under this approximation, the lowest-order modulatory term beyond the bilinear one is 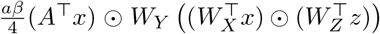, so we again expect the diagnostic to localize the violation to *x*.
- *Exponential:* Here, there are two lowest-order modulatory terms beyond bilinear, namely 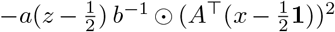 (nonlinear in *x* only) and 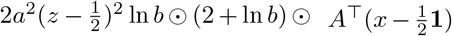 (nonlinear in *z* only), where 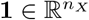 is the vector of 1s and 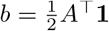. Thus, the diagnostic should detect a violation in both *x* and *z*, with more power in *x* for *a <* 1, since the *x* term is first order in *a* and the *z* term second order.

Across all generators, the diagnostic detected a violation of bilinearity and localized it to the expected factor (*x* or *z*). Under a bilinear generator, the diagnostic correctly reported no violation.

### B.12 Sample-efficiency of MICs compared to one-hidden-layer MLPs

In simulations of nonlinear generators (Appendix B.10), MICs correctly detected the geometry and presence of modulation, but with reduced sensitivity: the EV_mod_ captured by MICs was typically smaller than the ground-truth, especially for strong modulation (Fig. S5). We therefore asked how much more EV_mod_ a flexible nonlinear model could capture. Specifically, we compared MICs to a one-hidden-layer perceptron (MLP; ReLU activation function, *h* hidden units), a universal approximator of any continuous function [66]. We trained the MLP on exactly the same inputs and target output as the modulation term in MICs: it receives [**X**^*′*^, **Z**] and predicts **Y**^*′′*^, the residuals of the joint additive term in MICs. The MLP has *h* (*n*_*X*_ + *n*_*Y*_ + *n*_*Z*_) + *h* + *n*_*Y*_ parameters, which even at *h* = *m*_*C*_ are more than that of MICs’ modulation term (*m*_*C*_(*n*_*X*_ + *n*_*Y*_ + *n*_*Z*_)). We swept *h* ∈ {*m*_*C*_, 10*m*_*C*_, 100*m*_*C*_}, up to 100 times more parameters than MICs. We used mean-squared-error loss, Adam (learning rate 10^*−*3^, 200 epochs, minibatch size 32) and ridge regularization selected on held-out data (penalty grid 10^*−*5^ to 10^*−*1^). We quantified the MLP’s cross-validated explained variance 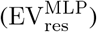 as the increment in explained variance over the joint additive model, exactly as we did for MICs EV_mod_.

Note that while EV_mod_ isolates the bilinear modulation term, 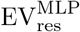 captures any nonlinear structure in the residuals, whether from modulation or from nonlinearity in the marginal *x* → *y* and *z* → *y* mappings. Thus, the residual explained variance from the MLP is an upper bound on the variance attributable to modulation of any form: 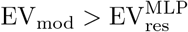 establishes that the flexible model does not extract more from the residuals than MICs does, whereas the reverse inequality is not specifically attributable to modulation.

Under a bilinear generator, we found 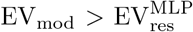 regardless of sample size *N* (Fig. S6a). Even under a strongly nonlinear generator (parallel-axis rotation with amplitude *a* = *π*; Appendix B.10), MICs fares better than the MLP at small *N*, though the MLP eventually overtakes it for large enough *N* (Fig. S6c). Under a milder nonlinearity (parallel-axis rotation with *a* = 2*π/*5), MICs exceeded MLP at all *N* considered and degraded much more slowly as *N* decreased (Fig. S6b). Thus, while the MLP has higher flexibility to capture nonlinear modulations, this comes at the cost of reduced sample efficiency and interpretability.

### B.13 Statistical testing of modulatory and AIS EV

#### Motivation

The 1-SEM rule used for rank selection (Algorithm 1; [16, 51]) is a heuristic that can yield spurious AIS or modulatory EV in a fraction of cases when the ground-truth *m*_*A*_ or *m*_*C*_ is zero. Across the 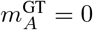 or 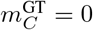 scenarios of Fig. S1 (*rand*, Step 1 on), the rule alone retained *m*_*A*_ ≥ 1 and *m*_*C*_ ≥ 1 in 18 % and 34 % of simulations, respectively. We therefore complement rank selection with a permutation test, applied hierarchically to AIS (*m*_*A*_ *>* 0) and to modulation (*m*_*C*_ *>* 0), that decides on held-out data whether each candidate term predicts *Y* better than expected under a null in which it is temporally unrelated to the test-set residuals it is meant to explain.

#### Test construction

We split the dataset into two non-overlapping halves (first and second half of samples). For each split, the MICs pipeline (rank selection, ridge regularization, and ALS) is fit on one half and evaluated on the other, so that all fitting and hyperparameter selection are confined to the training half. On the held-out half we form three predictions of *Y* of increasing complexity,

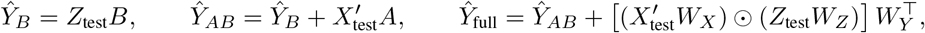

where *X*^*′*^ = *X* − *ZD* uses the regression *D* fit in Step 1 of Algorithm 1. The observed held-out EV increments

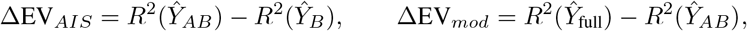

quantify the variance in *Y*_test_ captured by AIS and by modulation, respectively. *R*^2^(*Ŷ*) denotes the coefficient of determination of real *Y* from prediction *Ŷ* compared against *Y* training mean.

To test significance of ΔEV_*mod*_, we shuffle in time the predicted modulation component 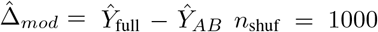 times. For time series, we apply random circular shifts uniformly drawn from [⌊*N*_test_*/*4⌋, ⌊3*N*_test_*/*4⌋], where *N*_test_ denotes the number of test samples. For i.i.d. samples from simulations, we randomly permute 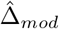 across all test samples. We then recompute 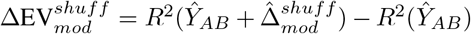 on each shuffle. Circular shifting destroys the temporal alignment between 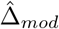 and the residual *Y*_test_ − *Ŷ*_*AB*_ while preserving the marginal distribution and autocorrelation of 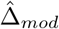, so that adding back a misaligned 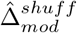 to *Ŷ*_*AB*_ generally inflates the sum of squared errors and yields a null statistic with negative mean. A genuine modulatory contribution should therefore exceed essentially all shuffles. Null EV are averaged across the two splits, and a one-tailed *p*-value is computed empirically, comparing ΔEV_*mod*_ to the ΔEV^*shuff*^ distribution. ΔEV_*AIS*_ significance is tested identically with 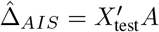 in place of 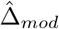 and *Ŷ*_*B*_ in place of *Ŷ*_*AB*_ . A non-zero rank is retained only if *p <* 0.05; otherwise the corresponding term is fixed to zero in all downstream analyses (geometry indices, constrained-fit comparisons).

Testing *m*_*A*_ and *m*_*C*_ with this procedure reduces the average false-positive rate at 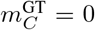 from 34 % to 0 % across the 16 scenarios of Fig. S1 while preserving a true-positive rate of 100 % at 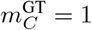 (vs. 100 % without the test); analogous values for *m*_*A*_: 18 %, 4 %, 100 %, 100 %.

### B.14 Details and further analyses of V1-V2 data

#### Experimental procedures and preprocessing

We re-analyzed the publicly available V1-V2 dataset of Zandvakili and Kohn https://crcns.org/data-sets/vc/v1v2-1, a subset of the recordings described in [55] comprising the largest and best retinotopically aligned populations, with V2 electrodes targeted to the input layers, previously analyzed by Semedo et al. [16] and by Gokcen et al.[56], to which we refer for full experimental details. Briefly, V1 (Utah array) and V2 (tetrodes) populations with aligned receptive fields were recorded simultaneously in anesthetized macaques viewing drifting gratings of 8 orientations *θ* (22.5^*°*^ steps; 300-400 trials per orientation, 1.28 s each). The dataset consists of 5 sessions from 3 animals (88-159 V1 and 24-37 V2 units per session). Following [16], we used 100 ms spike-count bins from 160 to 1160 ms after stimulus onset, retained units with firing rate ≥ 0.5 spikes/s, and subtracted for each unit, orientation and time bin the so called Peri-Stimulus-Time-Histogram, or PSTH, that is time-dependent firing rate averaged over trials with the same orientation, so that analyses target trial-to-trial residual co-fluctuations. Whereas [16] and [56] fit an independent model within each orientation, we fit a single model across all orientations, taking V1 units as source *X*, V2 units as target *Y*, and grating orientation as modulator *Z*. Because the PSTH is removed, orientation has no first-order effect on either population and the additive terms *B* and *D* explain zero variance; EV_mod_ thus quantifies whether the V1-to-V2 residual mapping depends on *θ*. We used two encodings of *θ*: a one-hot encoding of the eight orientations (*Z* ∈ ℝ^8^), which imposes no functional form on how the interaction depends on *θ*, and a periodic encoding *Z* = (sin 2*θ*, cos 2*θ*) ∈ ℝ^2^. Unless stated otherwise, results refer to the one-hot encoding.

**Figure S9:**
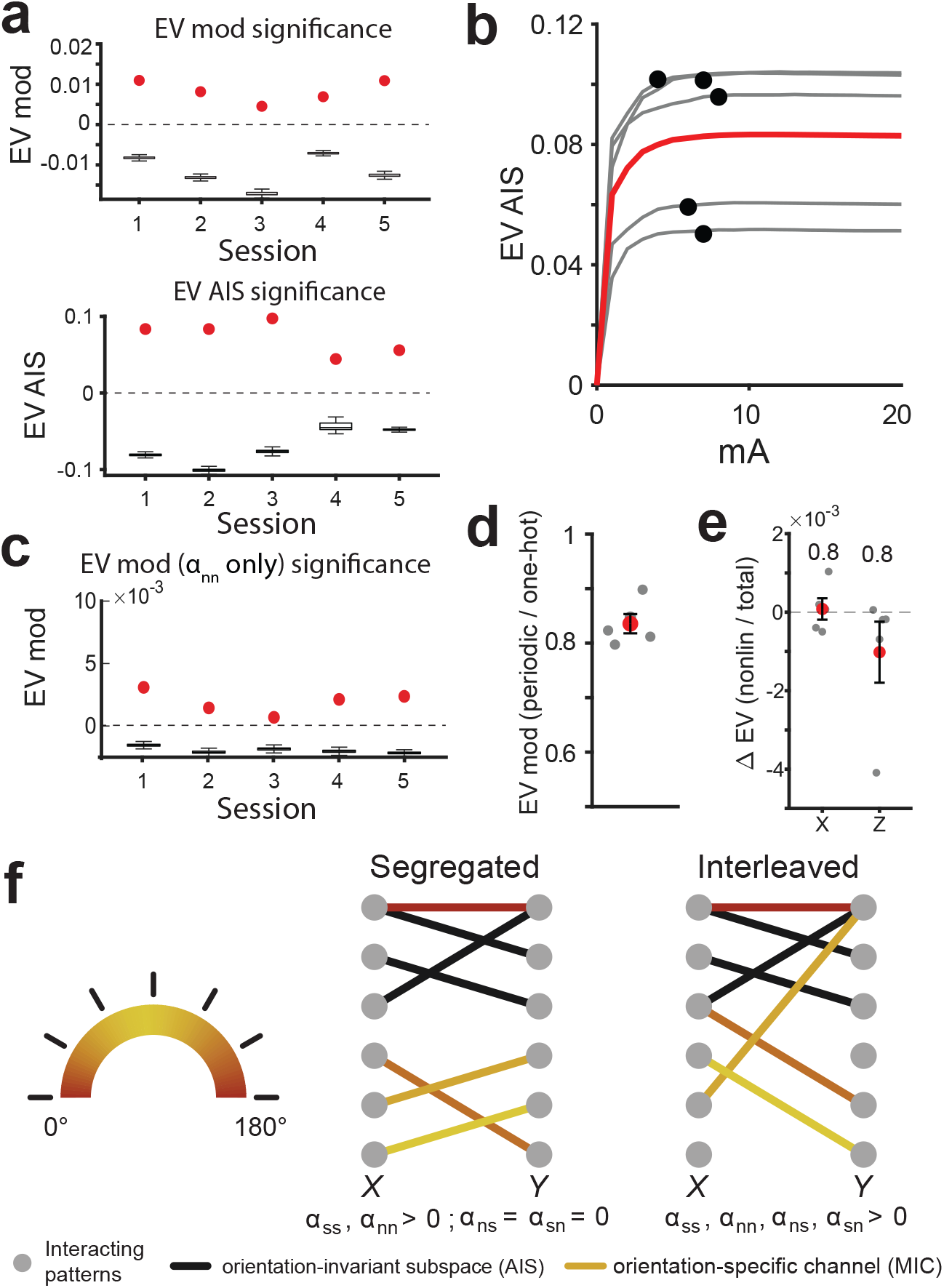
Additional analyses of V1-V2 benchmark data. All panels use the one-hot encoding of orientation unless stated. (a) Single-session significance of EV_mod_ (top) and EV_AIS_ (bottom) against the residual-shuffle null (Appendix B.13). Boxes: null median and IQR; whiskers: 1.5 × IQR; red dots: observed held-out EV. Both terms were significant in 5/5 sessions. (b) EV_AIS_ as a function of AIS rank *m*_*A*_; gray lines: individual sessions; black dots: single-session selected rank; red: across-sessions mean. (c) As (a, top), for MICs constrained to the *nn* component only (*w*_*X,i*_ orthogonal to the V1 AIS and *w*_*Y,i*_ orthogonal to the V2 AIS; Appendix B.8). Despite 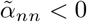 (Fig. 4d), *nn*-only modulation was significant in 5/5 sessions (EV_mod_ = (0.24 ± 0.05)%). (d) Ratio between EV_mod_ obtained with the periodic encoding *Z* = (sin 2*θ*, cos 2*θ*) and with the one-hot encoding; black dots: sessions; red: mean ± SEM ((83.6 ± 1.8)%). (e) Bilinearity diagnostic (Appendix B.11): change in cross-validated EV when higher-order terms in *x* or *z* are added to the fitted MICs (ΔEV_NL_). Gray dots: sessions; red: mean ± SEM; numbers: one-tailed binomial test *p*-values. (f) Schematic of segregated versus interleaved organization of orientation-specific interactions. Nodes: V1 (*X*) and V2 (*Y*) interacting activity patterns; black edges: *θ*-invariant interaction (AIS); colored edges: MICs, colored by the orientation at which they are most strongly engaged. Left) segregated: each MIC lies inside or outside the AIS at both ends (*ss, nn*), so *α*_*ns*_ = *α*_*sn*_ = 0. Right) interleaved: MICs also couple an AIS pattern on one side with a non-AIS pattern on the other (*ns, sn*), as observed in real data (Fig. 4d).

#### MICs fitting and main analyses

We fit the pipeline of Algorithm 1 separately to each session as for the ACA-VIP data (Appendix B.15): *K* = 10 temporally contiguous cross-validation folds, all variables z-scored at the fitting stage, and AIS and MICs ranks capped at 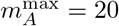 and 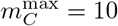 (mean selected ranks *m*_*A*_ = 4.8, *m*_*C*_ = 6.4). Single-session significance of EV_mod_ and EV_AIS_ was assessed via the residual-shuffle permutation test of Appendix B.13 with *n*_shuf_ = 1000 shuffles per session (Fig. S9a). In Fig. 4b, EV_mod_ is normalized by EV_AIS_ to allow comparison across datasets. For the example MIC of Fig. 4c, orientations were ordered by the rank of their modulator score *Zw*_*Z,i*_; the colorbar reports the angular difference of each orientation from the lowest-scoring one. The MIC correlation slope (Fig. 4c) was computed as for ACA-VIP data, except that the modulator score takes only 8 values: the Pearson correlation between *Xw*_*X,i*_ and *Y w*_*Y,i*_ was computed within each orientation and regressed against the orientation score rank-normalized to [0, 1]; slopes were averaged across MICs and then across sessions. Geometry indices (Fig. 4d, top) and constrained fits (Fig. 4d, bottom) were computed as for ACA-VIP data, with “V1” and “V2” denoting the constraint applied to the source and target factor, respectively; the *nn*-only model (Fig. S9c) constrains both factors to the orthogonal complement of the corresponding AIS (Appendix B.8). Because sessions from the same animal are not independent, statistics in Fig. 4d,h use linear mixed-effects models with a random intercept per animal. For each null-subtracted index, and for each selected constrained-fit comparison (Appendix B.8), we modeled the per-session value (the index, or the difference in EV_mod_ between the two fits) as *d* ∼ 1 + (1 | animal) and tested the intercept against zero (two-tailed; Holm corrected across the four indices and across the three comparisons of Fig. 4d, respectively). For simplicity, tests in the supplementary figures take sessions as the unit of analysis. To test whether the modulation is captured by a smooth periodic function of *θ*, we refit MICs with the periodic encoding, using the same pipeline and folds, and report per session the ratio of its cross-validated EV_mod_ to that of the one-hot encoding (Fig. S9d).

### B.15 Details and further analyses of ACA-VIP data

#### Experimental procedures and preprocessing

We collected simultaneous dual-color two-photon calcium imaging of ACA axonal projections in V1 and V1 VIP interneurons in head-fixed mice during viewing of visual stimuli, while continuously tracking pupil diameter and running speed. The dataset is publicly available at https://osf.io/2ekj4/. All procedures were approved by the Massachusetts Institute of Technology Committee on Animal Care and conformed to the National Institutes of Health guidelines for the care and use of laboratory animals. VIP-Cre mice (Jackson stock no: 031628, RRID:IMSR_JAX:031628) were anesthetized with isoflurane (1.5%) and given preemptive analgesia (extended release buprenex, 1mg/kg, and meloxicam, 5mg/kg, s.c.). After being placed in a stereotaxic frame and prepared for surgery, a small craniotomy was made above the ACA (AP: +1, ML: -0.3, DV: -0.9) and 0.3 ul of AAV1-hSyn-axon-GCaMP6s [67] was injected into the ACA with a micropipette. A larger craniotomy (3mm) was made above the V1 and 4 small injection of 0.15*µ*l AAV9-DIO-jRGECO1a [68] was made into the V1 before placing the cranial window and securing it on the skull with dental cement. Visual stimuli were drifting gratings (8 directions × 3 contrasts, 10 trials per condition; 240 trials/session) or natural movies (5 movies ×3 contrasts, 10 trials per condition; 150 trials/session). Moreover, 25 of 28 sessions included 15 mild airpuffs (compressed air at 40 psi for 0.3 s) per session in alternating trial blocks to evoke arousal events. Frames were acquired at 33 Hz and every other frame was averaged, yielding an effective sampling rate of ∼ 16.5 Hz. Imaging used a Prairie Ultima IV two-photon microscope with a MaiTai DeepSee laser (Spectra-Physics) tuned to 965 nm, resonant-galvo scanning (Prairie View 5.4) and a 20× objective (XLPlan N, 1.00 NA, Olympus) at 4× digital zoom. Behavioral apparatus, stimulus presentation and synchronization were as in [59]. Pupil diameter was extracted from 20 Hz face video by tracking 8 pupil-boundary markers with DeepLabCut. It was computed as the horizontal distance between two opposing markers, scaled by the median vertical-to-horizontal diameter ratio, and converted to mm (60.8 pixels/mm). Artifacts (frame-to-frame decreases below the 5th percentile of negative changes) were replaced by modified Akima interpolation, values ≥ 2.5 mm or missing were filled with the preceding value, and the trace was smoothed with a 1 s moving median. Running speed (cm/s) was computed from a rotary encoder (0.0158 cm/count), rectified, resampled to 20 Hz, and smoothed with a causal 0.5 s moving median. Calcium traces were neuropil-corrected as *F*_ROI_ = *F*_ROI,raw_ − 0.7 *F*_neuropil_. To obtain single-axon traces, ACA boutons were grouped into putative single-axon clusters by hierarchical clustering on pairwise temporal correlations, and within-cluster traces were averaged with weights proportional to ROI signal-to-noise ratio. All ACA-axon and VIP-neuron traces were linearly detrended within session prior to analysis. All variables were linearly interpolated onto a common 20 Hz time vector (∼ 24,600 samples/session, ∼ 20.5 min). We used 28 sessions (15 movies; 13 gratings) collected across 3 mice (1 male, 2 female; aged 3-5 months during imaging), taking as response variables the per-axon and per-neuron calcium traces. Included counts ranged 3-212 for ACA axons and 1-11 for VIP neurons. Wheel acquisition started slightly after and ended slightly before imaging; the time points without running speed (1–2% per session) were excluded.

**Figure S10:**
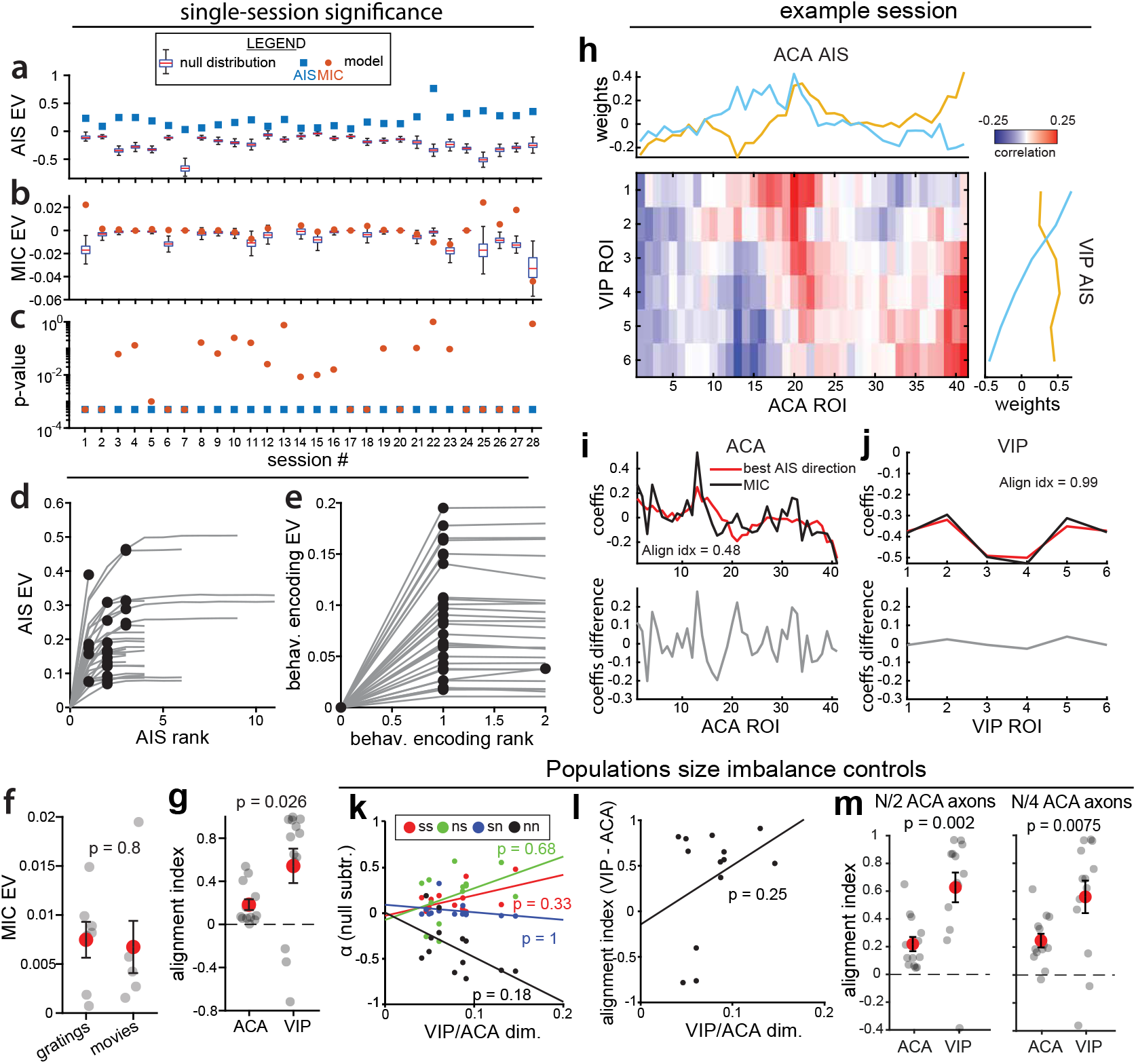
Additional analyses of ACA-VIP recordings. (a) Single-session AIS EV significance against a shuffle null (see App. B.13). Boxes: null distribution median and IQR; whiskers: 1.5 × IQR; dots: mean held-out AIS EV. (b) Same as (a) but for EV MIC. (c) Single-session empirical AIS EV and MIC EV p-values. (d) AIS EV as a function of AIS rank; gray lines, individual session; black dots, single-session selected rank. (e) Same as (d) for behavioral state encoding EV (term *B*). (f) Comparison of MIC EV in sessions with different visual stimuli (drifting gratings vs. natural movies). Gray circles: individual sessions; red circles: mean ± SEM across sessions. Only sessions with significant (shuffle test *p <* 0.05, see (a)) are shown. (g) MIC-AIS alignment index for ACA weights (left) and VIP weights (right) across sessions. (h) AIS low-rank decomposition for one example session (*m*_*A*_ = 2). Heatmap: ACA-VIP cross-correlation matrix after regressing out behavioral state (Step 1, Alg. 1). Top: ACA AIS weights. Right: VIP AIS weights. For visualization purposes, ACA and VIP ROIs were sorted using optimal leafs order of hierarchical clustering (Euclidean distance, Ward linkage). (i) MIC alignment with AIS for ACA (*m*_*C*_ = 1). Top: MIC weights (black) and their best approximation within the AIS (red). Bottom: residual difference. Alignment index reported in panel. (j) Same as (i) for VIP. (k) Null-subtracted geometry indices *α*_*ss*_ (red), *α*_*ns*_ (green), *α*_*sn*_ (blue), and *α*_*nn*_ (black) as a function of the VIP:ACA population size ratio. Lines: best linear fits. (l) Same as (k), but for the difference between VIP and ACA MIC-AIS alignment index. (m) Same as (g), after refitting MICs with ACA axons randomly subsampled to half (left) or a quarter (right) of each session’s population; per-session indices are averaged over 10 random subsamples. P-values show two-tailed t-test in (f), one-tailed paired t-test in (g,m), and Bonferroni-corrected Pearson’s correlation tests in (k,l).

**Figure S11:**
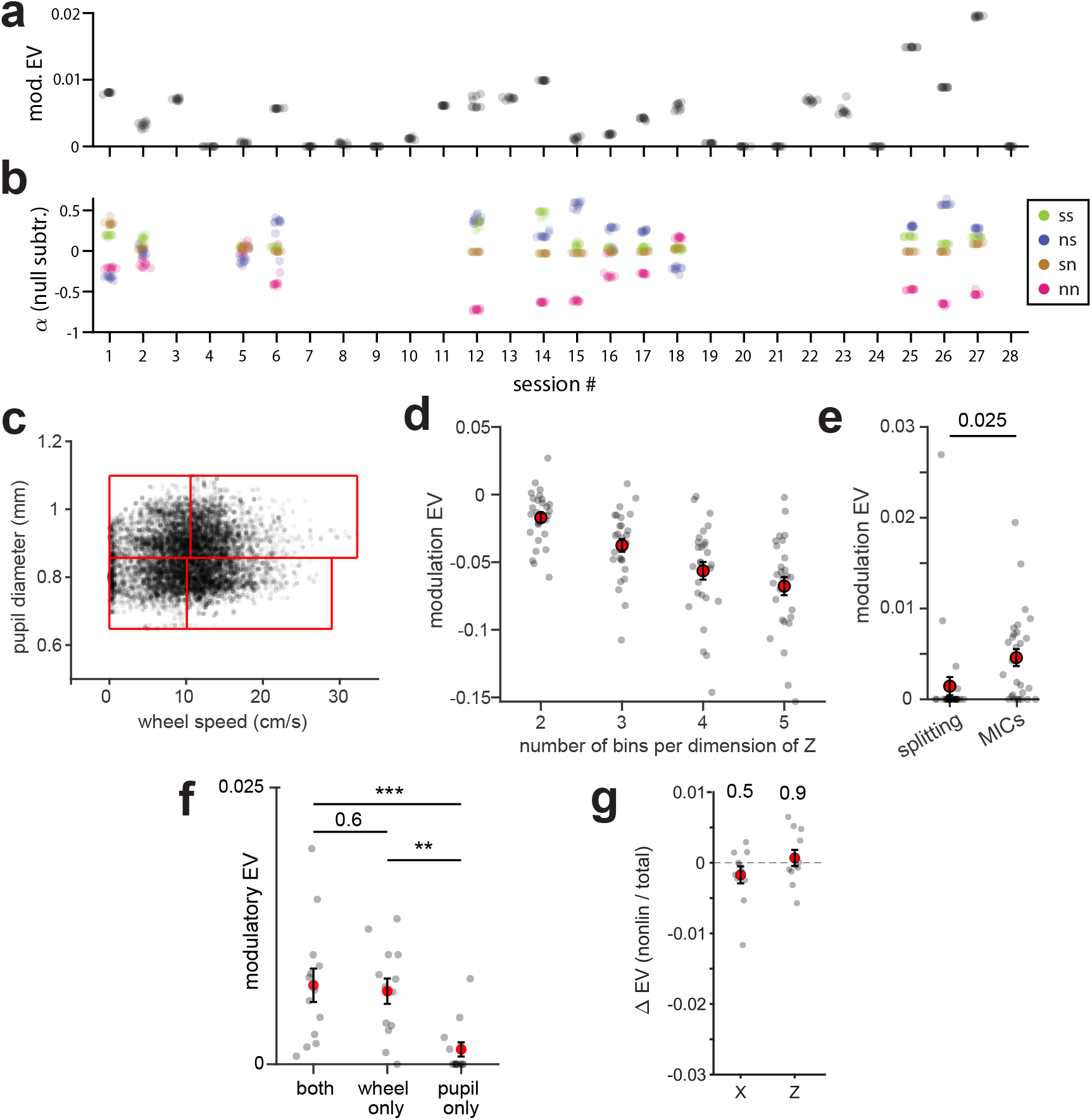
Robustness and control analyses of ACA-VIP recordings. For each of the 28 sessions, MICs were re-fitted 10 times with independently drawn random ALS initializations, holding all per-session ranks (*m*_*A*_, *m*_*B*_, *m*_*C*_, *m*_*D*_) fixed at the values selected in the main text. **(a)** EV_mod_ for each session and ALS seed (one dot per seed). **(b)** Null-subtracted geometry indices 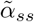 (green), 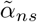(blue), 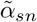 (brown), 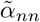(pink) for the 13 sessions with significant EV_mod_, shown for each seed; geometry was not interpreted for sessions without significant modulation. (c) Splitting-based alternative to MICs (Appendix B.15), example session: behavioral state is split into *N*_split_ wheel-speed quantile bins, each further split into *N*_split_ pupil-diameter quantile bins (red boxes; *N*_split_ = 2), and a separate AIS is fit within each bin. (d) Splitting-based EV_mod_ as a function of *N*_split_. (e) Splitting-based EV_mod_ at the best *N*_split_ per session versus MICs EV_mod_. (f) EV_mod_ with *Z* comprising both wheel speed and pupil diameter (both, as for all other analyses), wheel speed only (wheel only), or pupil diameter only (pupil only). (g) Bilinearity diagnostic, as in Fig. S9e. Gray dots: sessions (d, e: all 28; f, g: 13 with significant EV_mod_); red: mean ± SEM. P-values: two-tailed t-test in (e,f), one-tailed binomial test in (g). **, *p <* 0.01; ***, *p <* 0.001.

#### MICs fitting and main analyses

We fit the pipeline of Algorithm 1 separately to each session, with *K* = 10 temporally contiguous cross-validation folds to avoid leakage between training and test sets due to temporal autocorrelation. We took ACA axons as source *X* and VIP neurons as target *Y*, consistent with anatomical directionality, and defined the modulator *Z* ∈ ℝ ^2^ as the joint, within-session-standardized vector of pupil diameter and wheel running speed. AIS and MICs ranks were capped at 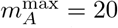 and 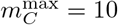 for computational efficiency. The MIC correlation slope (Fig.4g) was computed per channel by projecting *X, Y, Z* onto *w*_*X,i*_, *w*_*Y,i*_, *w*_*Z,i*_, partitioning *Zw*_*Z,i*_ into 5 equipopulated quantiles, computing the Pearson correlation between *Xw*_*X,i*_ and *Y w*_*Y,i*_ within each quantile, and taking the ordinary least squares slope of these five correlations against quantile index; slopes were averaged across MICs and then across sessions. Geometry was characterized via null-subtracted indices 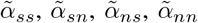 (Fig. 4h, top), where 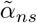 and 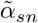 correspond to recruitment of private ACA (source) and VIP (target) dimensions, respectively. Each index was obtained by subtracting from the fitted *α* the mean over 1000 random rotations of *w*_*X,i*_, *w*_*Y,i*_ on their unit spheres (Appendix B.7.3). Constrained-fit EV_mod_ (Fig. 4h, bottom) further restricts *w*_*X*_ and/or *w*_*Y*_ to lie within the corresponding AIS following Appendix B.8, with “ACA” and “VIP” denoting the constraint applied to the source and target factor, respectively. Statistics for main text analyses were computed as for V1-V2 data (Appendix B.14), with Holm correction across the four constrained-fit comparisons of Fig. 4h. For simplicity, tests in the supplementary figures take sessions as the unit of analysis. Single-session significance of EV_mod_ and EV_*A*_ was assessed via the residual-shuffle permutation test of Appendix B.13 with *n*_shuf_ = 1000 shuffles per session (Fig. S10a-c). Across all 28 sessions, EV_mod_ correlated positively with the number of recorded VIP neurons (Pearson *r* = 0.64, *p* = 1.9 × 10^*−*4^), consistent with lower sensitivity in sessions with few target neurons.

##### Additional analyses

To visualize the MIC-AIS geometry in one example session (Fig. S10i), we computed the best approximation of each MIC factor within the corresponding AIS as *ŵ*_*X*_ = *P*_*X*_ *w*_*X*_ and *ŵ*_*Y*_ = *P*_*Y*_ *w*_*Y*_, with *P*_*X*_, *P*_*Y*_ the orthogonal projectors onto the source and target AIS. To compare alignment across sessions (Fig. S10g), we summarized per-session MIC-AIS alignment by a normalized index that rescales the observed squared cosine by chance alignment,

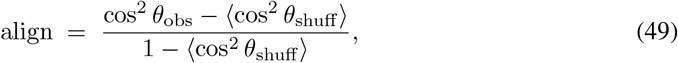

where cos^2^ *θ*_obs_ = ∥*P w*∥ ^2^*/* ∥*w*∥ ^2^ for the source (*w* = *w*_*X*_, *P* = *P*_*X*_) or target (*w* = *w*_*Y*_, *P* = *P*_*Y*_) factor (see Appendix B.7.2), and ⟨cos^2^ *θ*_shuff_⟩ is the same quantity averaged over uniform random samples of *w* on the unit sphere, matched in dimension and AIS rank. align = 0 corresponds to chance alignment given the dimensionality; align = 1 to a fitted MIC lying entirely within the AIS. We used a one-tailed test for this analysis under the hypothesis - motivated by geometry analyses in Fig 4 - that VIP MICs are more aligned to the AIS, compared to ACA. To rule out that the asymmetric alignment between ACA and VIP factors was driven by the typical *n*_ACA_ : *n*_VIP_ ≈ 15 population-size imbalance of sessions with significant EV_mod_, we regressed the per-session ACA-vs-VIP alignment difference and the four null-subtracted *α* indices against the ratio *n*_VIP_*/n*_ACA_. As a direct control, we refit MICs after randomly subsampling ACA axons to *N/*2 and *N/*4 of each session’s population (10 subsamples per session), and compared ACA and VIP alignment indices averaged across subsamples, as in Fig. S10g. VIP alignment remained significantly larger than ACA alignment, with a similar effect size, in both cases (Fig. S10m). Furthermore, to test whether airpuff-evoked arousal events drove the recovered modulation, we ran a control analysis recomputing EV_mod_ after excluding the 10 s window following each airpuff onset (not shown); EV_mod_ did not differ significantly from the full-session value, suggesting that modulation is driven by endogenous fluctuations in behavioral state rather than puff-evoked transients. Consistently, 2 of the 3 puff-free sessions were among the 13 sessions with significant EV_mod_. Finally, to assess robustness of MIC fits to the non-convexity of the ALS objective (Appendix B.5), we re-fitted MICs on each session with 10 independent random ALS initializations, holding the per-session ranks (*m*_*A*_, *m*_*B*_, *m*_*C*_, *m*_*D*_) fixed at the values selected in the main text. For each seed we recomputed EV_mod_ for all sessions and the null-subtracted *α* indices for the 13 sessions with significant EV_mod_ (Fig. S11). Both EV_mod_ and the null-subtracted *α* indices were tightly clustered across seeds, indicating that the ACA-VIP findings are stable under the non-convexity of the ALS objective.

As an alternative to fitting MICs to the ACA-VIP dataset, we split behavioral state into bins and fitted a separate ACA-VIP interaction model in each bin (Fig. S11c-e). Specifically, we first split into *N*_split_ bins by wheel speed quantile, and then further split into *N*_split_ sub-bins by pupil diameter quantile, for a total of 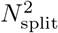 bins. We then followed the same fitting procedure as in Fig. S4 to obtain a splitting-based EV_mod_ value directly comparable to that outputted by MICs. Selecting *N*_split_ in each session to maximize EV_mod_ resulted in (mean ± SEM across session) EV_mod_ = 0.0018 ± 0.0010 (compared to EV_mod_ = 0.0046 ± 0.0009 for MICs across all sessions; *p* = 0.03, two-tailed t-test). The same temporally contiguous cross-validation folds were used for fitting and assessing the performance of MICs and the splitting alternative. Repeating the same procedure with the order of wheel and pupil reversed when selecting the bins for *Z* yielded splitting EV_mod_ = 0.0014 ± 0.0010 (*p* = 0.02, two-tailed t-test comparing to MICs EV_mod_). In sum, increasing the number of splits systematically decreased the performance of the aggregate model, resulting in negative values of EV_mod_, confirming that the binning approach is sample-inefficient.

To identify which behavioral variable drives the modulation, we refit MICs with *Z* comprising wheel speed only or pupil diameter only (Fig. S11f). Dropping pupil diameter did not reduce EV_mod_ (0.0066 ± 0.0011 wheel only vs. 0.0071 ± 0.0012 both, *p* = 0.64), whereas dropping wheel speed did (0.0013 ± 0.0006 pupil only; *p* = 0.0007 vs. both, *p* = 0.003 vs. wheel only).

### B.16 Computational resources

All simulations and real data analyses ran on a workstation with a 24 cores Intel i9-14900K processor and 192 GB of RAM, running Windows 11 and MATLAB R2025b, with parallelization over 20 workers (MATLAB Parallel Computing Toolbox). The pipeline additionally requires the Statistics and Machine Learning Toolbox.

Simulations in Figure 2a-e ran in approximately 10 minutes. All other simulations in Figures 2, 3, and S1–S8 ran in less than one day. Real data analyses in Figures 4 and S9–S11 also completed in under one day on the same machine, except for the residual-permutation test and the constrained model fitting, which took approximately two days.

Code implementing the MICs pipeline, the simulations, and the real-data analyses is available at https://github.com/mcelotto/multiplicative_interaction_channels. The full ACA-VIP dual-color two-photon imaging dataset is available at https://osf.io/2ekj4/.

## Footnotes

1 Unlike the RRR subspace - identifiable only as a column space - the CP components of the resulting tensor *C* satisfying Kruskal’s condition [49] are individually identifiable (Appendix B.3); hence we call them *channels*, not subspaces.

